# Breaking Through Microbial Defenses – Organic Acid-Based Deep Eutectic Solvents as a Neoteric Strategy in Bacterial Biofilm, Persister, and Fungal Control

**DOI:** 10.1101/2025.03.25.645211

**Authors:** Tomasz Swebocki, Karolina Cieminska, Clovis Bortolus, Jérôme Muchembled, Meroua S. Mechouche, Justine Jacquin, Rabah Boukherroub, Kamel Haddadi, Ali Siah, Boualem Sendid, Aleksandra M. Kocot, Magdalena Plotka

## Abstract

This study explores the adaptation of organic acid-based deep eutectic solvents (OA-DESs) as effective antimicrobial agents. Having already demonstrated their efficacy against planktonic bacteria in our previous research, we now investigate their impact on more complex microbial forms, including biofilms, persister cells, and fungi (both human pathogenic and phytopathogenic). Our experiments revealed that OA-DESs effectively eradicated MRSA and *Escherichia coli* biofilms, inducing significant morphological changes. Notably, a three-log-unit reduction was observed for most OA-DESs at concentrations below 1% v/v, a remarkable achievement for this class of materials. Additionally, with only one exception, OA-DESs did not promote persister cell formation, underscoring their potential for complete biofilm eradication. In another part of our study, OA-DESs were compared to conventional DESs against *Candida albicans*, *Candida auris*, and *Aspergillus fumigatus*. Results showed that while individual DES components exhibited minimal activity, their combination effectively inhibited fungal growth and induced substantial morphological changes. Lastly, OA-DESs were tested against the phytopathogens *Zymoseptoria tritici* and *Venturia inaequalis*. Though their activity was less pronounced compared to pathogenic strains, most OA-DESs inhibited the growth of both fungi at the highest tested concentrations. Despite the broad scope of this study, we believe this study provides valuable insights into the potential of DESs as antimicrobial agents, offering a strong foundation for future research and innovation in this field.

## 1. Introduction

Microorganisms play a pivotal role in the environment – both in natural as well as in artificial, man-made. They can be found in almost every ecosystem on the Earth, from deep-sea hydrothermal vents (Zhou *et al*. 2022) to the human digestive system(McCallum and Tropini 2024). These tiny organisms, which include bacteria, fungi, viruses, and protozoa, contribute to a wide range of processes that are essential for environmental health and human well-being. These cover, among others, nutrient cycling (such as nitrogen and carbon cycles)(Arrigo 2005), bioremediation (Kumar and Bharadvaja 2020), food production(Sharma *et al*. 2020), waste treatment (Kumar, Sehgal and Gupta 2023), and biotechnology(Demain 2000) – with many of these fields often overlapping. Most of the microorganisms are harmless to humans, however, still many of them pose a threat to our health, either directly by causing infection or indirectly, e.g. by spoiling foods, which consumption may cause foodborne diseases. One way to counteract the unwanted effect of microorganisms on our health is a strict control of microbial population. This covers both the enhancement of the population of certain strains (e.g. via use of probiotics that target gut microbiota) as well as prevention of the contact with pathogens (e.g. use of antimicrobial agents). However, as easy as it may seem, the control of microbial population still remains a global challenge and is one of the most important aspects in the One Health strategy(McEwen and Collignon 2018a).

Over the years, many antibacterial strategies have been studied, developed and implemented. Some being straightforward – such as the use of extreme heat and pressure(Wuytack, Diels and Michiels 2002), UV irradiation(Coohill and Sagripanti 2008) or oxidizing agents(Miller and Britigan 1997). However, these methods, despite being effective are, due to their nature, used mainly in the circumstances where an aggressive and fast eradication is desired such as industrial surfaces, sterilization of labware and medical tools and sanitary upkeep. In more complex approaches, such as human, animal and plant treatment, a much more careful and selective approach is needed – mainly in order to protect healthy cells and tissues. Here, substances like antibiotics/antifungals or antimicrobial peptides are of great use.

The discovery of the first antibiotic (penicillin) by Alexander Fleming in 1928 revolutionized medicine and the approach to combating bacterial infections.(Uddin *et al*. 2021) Subsequently, more antibiotics were discovered and over time they began to play a key role in treating bacterial infections in humans.(Hutchings, Truman and Wilkinson 2019) Antibiotics affect cellular structures or metabolic processes, inhibiting the growth and cell division of bacteria, and their effects can be differentiated into bactericidal (cell death) and bacteriostatic (growth inhibiting).(Lobritz *et al*. 2015) In the context of antibiotic therapies, a critical point in maintaining public health is their proper use. The overuse of antibiotics in human medicine is a major contributor to the development of antibiotic resistance.(McEwen and Collignon 2018b; Uddin *et al*. 2021) Similar observations apply to veterinary medicine, where antibiotics are used both to treat and prevent bacterial infections among animals. Excessive use of antibiotics in animal agriculture also contributes to antibiotic resistance, which has consequences not only for animals (failure of antibiotic treatments in the future), but also affects human health through the food chain.(Hu *et al*. 2023) Antimicrobials are also used in agriculture to protect plants against bacterial and fungal diseases. However, the protection is mainly based on the use of chemical fungicides and the use of antibiotics in this sector is limited due to their impact on the environment and the risk of resistance emerging along the food chain and in environmental fungal strains able to spread in clinical wards, e.g. *Aspergillus fumigatus*.(Chowdhary and Meis 2018; Manyi-Loh *et al*. 2018; Hu *et al*. 2023) Nevertheless, antimicrobial resistance, including fungicide resistance in both agriculture and medicine, is a global threat to a public health. Excessive and inappropriate use of antibiotics in medicine, veterinary and agriculture contributes to the faster development of resistance, which threatens the health of humans, animals, plants and the environment. For this reason, alternative methods of fighting microorganisms are being sought. In this context, the following are used, among others: antibacterial peptides(Rima *et al*. 2021), phage therapy(Hatfull, Dedrick and Schooley 2022), probiotics(Ali *et al*. 2023), enzybiotics(Dams and Briers 2019), phytochemicals(AlSheikh *et al*. 2020; Khare *et al*. 2021), immunotherapy(Ramamurthy *et al*. 2021), laser and photodynamic therapies(Hamblin and Hasan 2004), quorum sensing inhibitors(Brackman *et al*. 2011) and nanotechnology(*Metal-Based Nanoparticles as Antimicrobial Agents: An Overview*).

Over the recent years, a new field of research emerged, related to the antibacterial properties of deep eutectic solvents (DESs). DESs are mixtures of two (binary) or more (ternary, tertiary…) substances that exhibit a eutectic point – liquefying each other and resulting in their new properties(Abbott *et al*. 2004). In these systems, one of the substances acts as a donor and the other as an acceptor of hydrogen bonds. Consequently, the mixture of these substances has a lower melting point than the parent substances themselves. Most of the research on DESs to date has focused on their use in green chemistry as they are an alternative to volatile or organic solvents. They were also considered as a drug delivery vehicle in (bio)medicine.(Javed *et al*. 2024) Lately, the number of reports on DESs in the context of combating bacterial infections is increasing. For instance, Akbar et al.(Akbar *et al*. 2023) evaluated the antibacterial potential of DESs against Gram-positive and Gram-negative bacteria. They showed that the strongest antibacterial activity was demonstrated by DESs based on methyltrioctylammonium chloride and glycerin as well as methyltrioctylammonium chloride and fructose. Both DESs showed strong activity against Gram-negative bacteria – *Pseudomonas aeruginosa* and *E. coli* with 40% and 68%, and 65% and 61% effectiveness, respectively. In turn, against Gram-positive bacteria, the strongest effect was recorded against *Bacillus cereus* and *Streptococcus pneumoniae* with an effectiveness of 75% and 51%, and 70% and 50%, respectively. In another study, Silva et al. investigated the potential of DESs containing fatty acids – capric, myristic, lauric and stearic acid to combat bacterial infections.(Silva *et al*. 2019) It was shown that DESs had a stronger antibacterial effect compared to isolated fatty acids, which indicates the synergistic effect of the components, and this could be observed particularly intensely in the DES composition of capric acid and myristic acid. All tested DESs showed activity against Gram-positive bacteria and fungi represented by *Candida albicans*, and the strongest antimicrobial effect was observed for DES with the composition capric acid and lauric acid. Therefore, DES based on these fatty acids were then used to assess biofilm removal. The effect of removing or detaching biofilms formed by Gram-positive bacteria, *C. albicans* and *E. coli* has been demonstrated. These results indicate a high potential for using DES based on capric acid and lauric acid to combat pathogenic microorganisms in medicine. In another study, Nysted et al.(Nystedt *et al*. 2023) investigated the effects of DESs based on choline chloride and xylitol, choline chloride and glycerol as well as betaine and sucrose for the elimination of *Staphylococcus aureus* and *P. aeruginosa* biofilms. Both choline and xylitol and choline chloride and glycerol DESs showed 4-6 orders reduction of viable cells in biofilm and 27-67% and 34-49% biomass removal for *S. aureus* and *P. aeruginosa*, respectively. Additionally, it was shown that these DESs inhibited the formation of biofilms of both strains at concentrations lower than or equal to 0.5×MIC, thus emphasizing the high potential for therapeutic use. These few examples show that DESs are an attractive alternative to antibiotics and more and more extensive research is justified on the possibility of their real use to fight microbial infections in times of global antibiotic crisis. The growing resistance of microorganisms to antibacterial substances is not the only factor determining and limiting the effectiveness of their use. An additional criterion limiting the success of therapy is also the form of occurrence of microorganisms – an organized group of microorganisms, i.e. biofilm, is more resistant to environmental factors and antibacterial substances than planktonic cells. Moreover, bacteria can isolate a subpopulation of persister cells, which in the presence of an antibiotic go into a dormant state or reduce their metabolic activity, and after cessation of exposure to the antibiotic, they regain full life functions. Importantly, persister cells differ from sensitive cells only phenotypically. In addition to bacterial infections, medicine and agriculture also struggle with fungal diseases. The difficulty of combating fungal infections is largely due to the much smaller spectrum of antifungal substances than in the case of antibacterial agents(Roemer and Krysan 2014), the occurrence of resistance to available antifungal agents(Perlin, Rautemaa-Richardson and Alastruey-Izquierdo 2017), the formation of biofilms(Fanning and Mitchell 2012), patients with impaired immune systems(Pfaller and Diekema 2007), the location of fungal lesions in the body(Casadevall and Pirofski 2003), slow growth(Perfect *et al*. 2001), and above all – similarity between fungal and human cells (eukaryotic structure), which makes it difficult to obtain substances selectively acting only against fungi(Brown *et al*. 2012). Therefore, in this study we propose the use of selected DESs, represented by organic acid-based DESs (OA-DES), including glycoline, T-glycoline, maloline, maline, and oxaline, to combat the most persistent and most difficult to eliminate forms of microorganisms – biofilms, persister cells and fungi. This work serves also as a direct continuation of our previous work being proof-of-concept, where these DESs were tested against planktonic bacteria(Swebocki *et al*. 2024). Moreover, the antifungal activity of these molecules against major plant pathogens such as *Venturia inaequalis* and *Zymoseptoria tritici,* responsible for apple scab and Septoria tritici blotch of wheat, respectively, will also be examined as a proof-of-concept for further studies.

## 2. Materials and methods

### 2.1. Chemicals and materials

All the organic acid-based deep eutectic solvents (OA-DESs) were synthesized from choline chloride or tetrabultylammonium chloride (TBACl) as well as organic acids: malic (MalA), malonic (MaloA), glycolic (GlycA), and oxalic (OxA). The non-OA-DESs were synthesized using glycerol (GLY), urea (U), choline chloride (ChCl), L-arginine (Arg), and ethylene glycol (EG). All the chemicals were supplied by Merck^®^ and were characterized by the purity of 98% and above.

### 3.2. Culture media

Culture media used for bacterial assessments were tryptic soy broth (TSB) for culturing of biofilms and persister cells assay, Mueller-Hinton agar (MHA) for colony counting, RPMI for antifungal assessment. All culture media were prepared according to the manufacturer preparation guide. The *Candida* strains and *A. fumigatus* were cultured on Sabouraud Medium. *V. inaequalis* and *Z. tritici* were cultured on Malt Extract agar and Potata Dextrose agar medium respectively. Media were supplied by Beckton Dickinson^®^ and Merck^®^.

### 2.3. Microbial strains

The following strains have been investigated in this work:

### 2.4. Methodology

All the microbiological manipulations were done in BSL2-class laboratory in the biosafety cabinet (class II).

#### 2.4.1. Synthesis of DESs

The selection and synthesis of DESs followed the procedure described in our previous work(Swebocki *et al*. 2024). Briefly, selected parent substances (PSs) were mixed in 1:2 ratio (ChCl or TBACl: organic acid), followed by gradual heating in dry block, with occasional vortex stirring to ensure homogeneity. Once the content of the tubes melted completely, the tubes were left to cool down, and a 3D printed insert with drying agent (silica gel with indicator, Merck^®^) was placed between the cup and the glass part of the tube, to ensure dry atmosphere. That prepared OA-DESs were then kept in the drawer at room temperature until further use.

#### 2.4.2. Biofilm culture

A single colony of MRSA or *E. coli* was transferred to 3 mL of tryptic soy broth (TSB) and incubated at 35°C until reaching OD_600_ of 0.5-0.6. The culture was then centrifuged (5 000 × g) for 5 min., followed by removal of supernatant. Then, bacteria were washed twice by resuspension of the pellet in 3 mL of potassium phosphate buffer (PPB), followed by centrifugation. The resuspended bacteria were then diluted to OD_600_ 0.01. Exactly, 1 mL of that prepared bacteria solution was transferred to a 24-well microtiter plates and left to adhere at room temperature for 2 h. After that time, liquid content of the wells was removed, and 1 mL of TSB was added to initiate the growth of the biofilm. Prior to incubation, the microtiter plates were covered with semi-permeable film, after which plates were put in the incubator at 35°C. The maturation of the biofilm lasted 24 h for MRSA with no medium change and 48 h for *E. coli* with a single medium change after 24 h. After maturation, biofilms were washed twice with 1 mL of PPB, and then treated with 1 mL of DES, dissolved in PBS for 4 h at 35°C. Finally, biofilms were washed three times with 1 mL of PPB to remove any remaining treating substance and medium.

#### 2.4.3. Minimum biofilm eradication concentration (MBEC)

The matured and treated biofilms were homogenized by scratching from the bottom surface of the well using pipette tip. The contents of the wells were then transferred into Eppendorf-type microtubes, vortex shaken and transferred onto 96-well microtiter plate with subsequent serial dilution using PPB. Next, the contents of the microtiter plate were plated onto MHA plates and incubated at 35°C overnight, after which colony counting was performed. The results were expressed as log(CFU/cm^2^) and two values were estimated based on the observations: MBEC (concentration at which we observe no colonies present) and MBEC99.9 (concentration at which a three-log-unit reduction of microbial population occurred).

#### 2.4.4. Live/Dead assay (LD)

Live/Dead assay was done according to a protocol published in the work of Sviridova et al.(Sviridova *et al*. 2022) In brief, PI and SYTO 9 were used to stain the matured biofilms before and after treatment. To this end, a premade solutions of these dyes were mixed in PBS (1 µL of SYTO 9 and PI stock solutions in 1 mL of PBS). After the biofilms were washed, 200 µL of solution of mixed dyes were added gently onto the surface of the biofilms and kept in dark for two minutes. After that time, the remaining solution was removed and the biofilms were imaged by means of brightfield microscopy using BioTek Cytation 5 Cell Imaging Multimode Reader (Agilent) equipped with 2.5×, 4× and 20× fold magnification lens system from Olympus in fluorescence mode.

#### 2.4.5. Persister cells assay (PC)

*E. coli* was tested with oxaline, maloline, T-glycoline, glycoline, and maline, while MRSA was tested with oxaline, maloline, T-glycoline, and glycoline. Overnight MRSA and *E. coli* cultures were refreshed in Miller-Hinton Broth (MHB) at a 1:100 dilution and incubated at 37°C until reaching an OD_600_ of 0.2. DESs were prepared in 2×MBEC in MHB, and bacterial cultures were added to the prepared DESs to achieve a final OD_600_ of 0.1 and 1×MBEC.

Bacterial solutions were incubated at 37°C with shaking, and samples were collected at intervals: every 5 minutes from 0 to 30 minutes, followed by 45 minutes, 1 hour, 2 hours, 3 hours, 6 hours, 24 hours, and 48 hours. Each sample was centrifuged at 5000 × g for 1 minute at room temperature, the supernatant was removed, and the pellet was resuspended in PBS and vortexed. The samples were then transferred to 96-well microtiter plates, serially diluted with PBS, and plated onto Luria agar (LA) plates. After overnight incubation at 37°C, colonies were counted, and the results were expressed as log CFU/mL.

#### 2.4.6. Antifungal assay

A single fungal colony was added to 15 mL of liquid Sabouraud medium and incubated overnight at 37°C with shaking, Yeasts were then centrifuged for 3 minutes at 600 × g, then washed with PBS (1X) and diluted with fresh Sabouraud medium to obtain a 0.5 MacFarland culture. Twenty µL of this culture was then added to 11 mL of Roswell Park Memorial Institute (RPMI) medium. One hundred µL of that prepared culture, was then added to a 96-well plate, together with 10 µL of the substance to obtain the desired concentration. The plate was then incubated at 37°C for 24 h, followed by reading at 600 nm using a spectrophotometer (FLUOstar Omega, BMG LABTECH).

#### 2.4.7. Phytopathogenic antifungal assay

Antifungal assays of OA-DES and non OA-DES against *V. inaequalis* and *Z. tritici* were performed on solid medium in 12-well plates. Malt agar and potato dextrose agar were used as mediums for *V. inaequalis* and *Z. tritici*, respectively. Four wells were used as reparations for each condition. The molecules were added to the culture medium after autoclaving, at 40 °C. In total, three concentrations (0, 0.1%, and 1%) of each molecule were used for each pathogen species. Plate inoculation was carried out by depositing, on the center of each well, 5 µL of fungal suspension at 10^5^ spores per milliliter of sterile water. Fungal spores were produced by cultivating the pathogen strains during 20 days for *V. inaequalis* and five days for *Z. tritici*. Antifungal activity was assessed by measuring the perpendicular diameter of each colony after 14 and 10 days of incubation at 20 °C for *V. inaequalis* and *Z. tritici*, respectively.

#### 2.4.8. Software

All the calculations and graphs were done using OriginPro 2025. AI tools implemented cover: Gramarrly, DeepL, Copilot and Google Translate – their use was limited to the style, punctuation and grammar control. Any AI-generated text was validated and modified by the authors. No data was generated by AI.

## 3. Results and discussion

### 3.1. Biofilm eradication efficiency of OA-DESs

#### 3.1.1. Minimum Biofilm Eradication Concentration (MBEC and MBEC99.9)

In our previous work,(Swebocki *et al*. 2024) we have investigated the antibacterial activity of various DESs. Our main conclusion was that OA-DESs displayed greater effectiveness when compared to conventional DESs. Moreover, we have also proved that DESs presented much greater potency when compared to their parent substances (PSs), highlighting the role of the vast network of hydrogen bonds, formed between PSs, on the antibacterial activity of the material. The mentioned study focused solely on the planktonic bacteria, without considering much more complex forms, which are responsible for great number of health and industry-related problems, that is – biofilms. With that in mind, we have decided to delve into the application of OA-DESs as effective biofilm eradication agents.

The first experiment carried out was the minimum biofilm eradication concentration (MBEC) assessment. Here, matured MRSA and *E. coli* biofilms were subjected to various concentrations of OA-DESs (glycoline, T-glycoline, maloline, maline, and oxaline). After being in contact for four hours, the biofilm was homogenized, diluted and plated on the agar Petri dishes in order to estimate the extent of the eradication of biofilm-enveloped bacteria.

**Figure 1.**
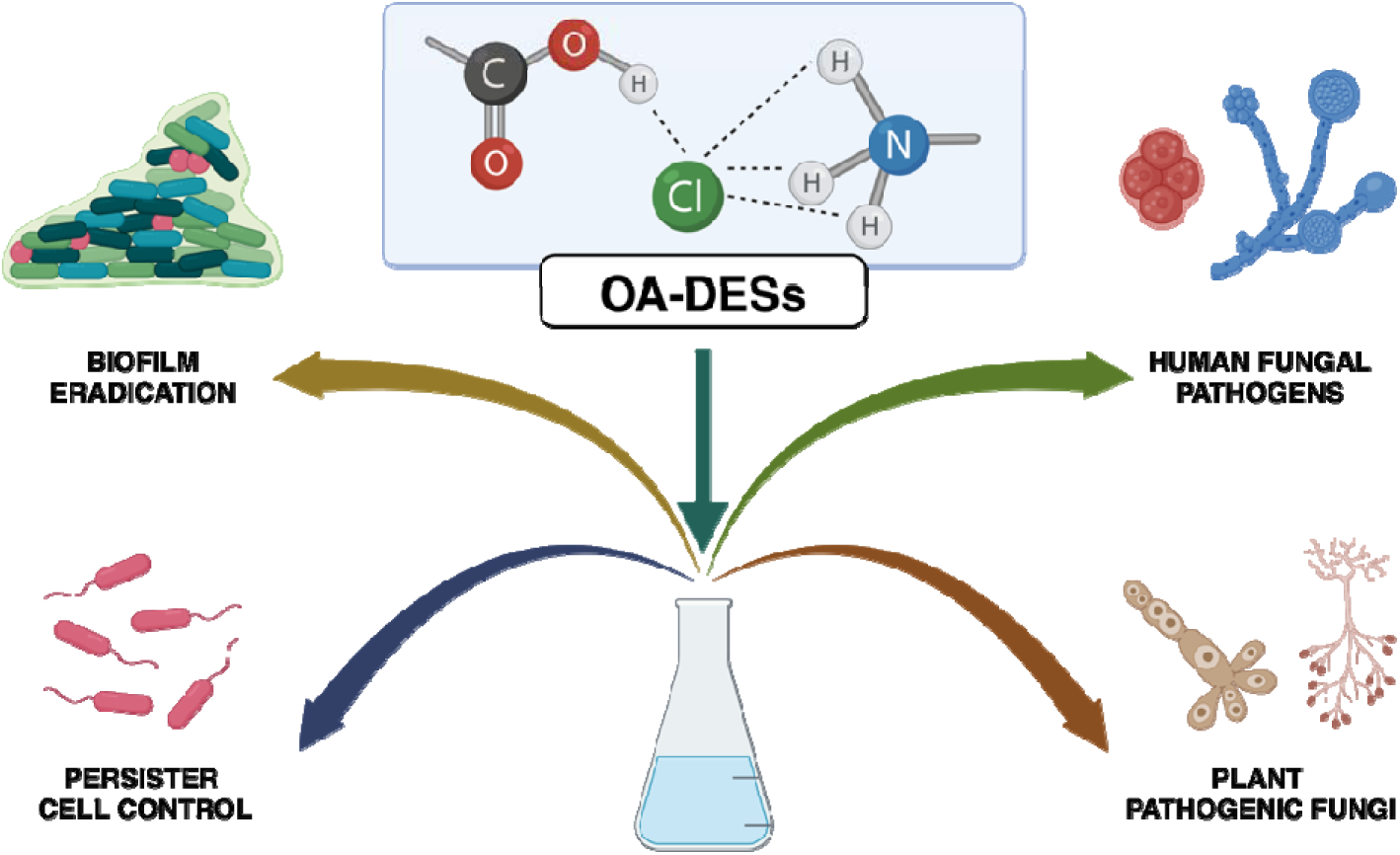
The objective of the present study – assessment of antimicrobial properties of OA-DESs targeting four main areas of interest: biofilm eradication, prevention of the emergence of persister cells, inhibition of the growth of fungi (both human and plant).

As seen in the **Figures 2**, **S1**, and **S2** and **Table 1**, in all but one cases, OA-DESs were able to effectively eradicate both types of the biofilms. When comparing the effectiveness of the investigated OA-DESs, it becomes clear that, in most cases, the activity of the DESs varied from substance to substance. The overall highest values of MBEC and MBEC99.9 (a three-log-unit drop of the population) were achieved for maline, with the second least effective DES being maloline, followed by glycoline, oxaline and finally the most effective OA-DES – T-glycoline. While the differences in MBEC and MBEC99.9 values between DESs provide us a general information on the antibacterial potency of the investigated substances, similarly to the previous studies of these compounds(Swebocki *et al*. 2024), one has to do a comparison between the strains, where the differences are much more pronounced.

**Figure 2.**
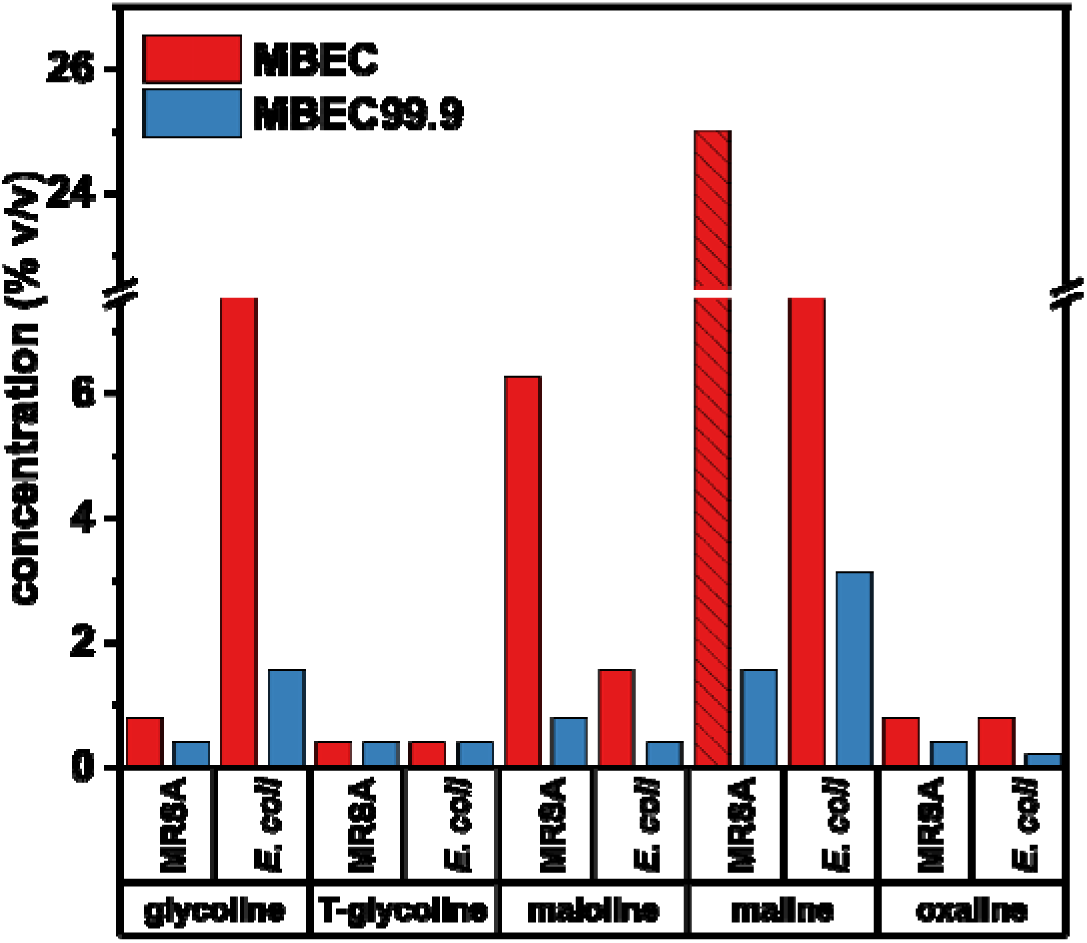
The MBEC assay of the investigated OA-DESs against Gram-positive (MRSA) and Gram-negative (*E. coli*) biofilms. The line pattern indicates that MBEC was not reached at the maximum concentration.

**Table 1.**
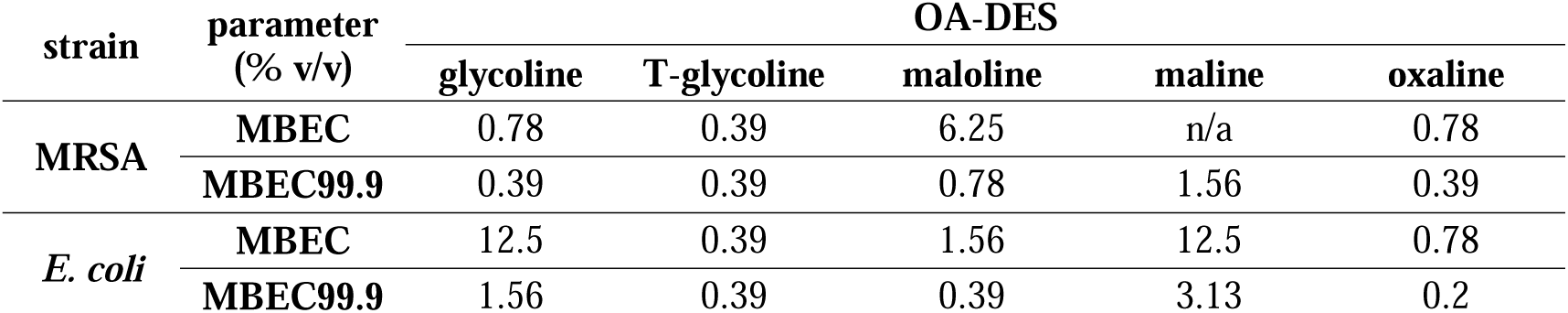
MBEC and MBEC99.9 values determined from figure 2.

**Table 2.** Strains of microbial organisms used in this study.

| name | identifier | family |
| --- | --- | --- |
| methicillin-resistant *Staphylococcus aureus* (MRSA) | ATCC® 43300 | bacteria (Gram +) |
| *Escherichia coli* | K-12 | bacteria (Gram –) |
| *Candida albicans* | ATCC® SC5314 | yeast |
| *Candida auris* | CHU 1.4.4.1.69 |  |
| *Aspergillus fumigatus* | ATCC® 204305 | yeast (mold) |
| *Zymoseptoria tritici* | T02596 | fungi (phytopathogenic) |
| *Venturia inaequalis* | Rs552 |  |

For glycoline, *E. coli* survivability was maintained up to 12.5% v/v, with an MBEC99.9 observed at 1.56% v/v. This OA-DES eradicated all MRSA cells at 0.78% v/v, with a significant population reduction at 0.39% v/v, making it uniquely more effective against Gram-positive species. For T-glycoline, both MBEC and MBEC99.9 were identical at 0.39% v/v for both strains, demonstrating consistent potency and no differentiation between strains. Maloline, the second least effective DES, showed higher MBEC values for MRSA (6.25% v/v) and *E. coli* (1.56% v/v), though MBEC99.9 values were notably lower, at 0.78% v/v for MRSA and 0.39% v/v for *E. coli* – showing higher antibacterial potency against Gram-negative species. Maline, the least effective OA-DES, displayed very high MBEC values (12.5% v/v for *E. coli*) and the highest MBEC99.9 value (3.13%). It was ineffective against MRSA biofilms even at 25% v/v, though a three-log-unit reduction occurred at 1.56% v/v – proving a similar higher affinity towards Gram-negative strain. Lastly, oxaline demonstrated high efficacy, with all MBEC and MBEC99.9 values below 1% v/v. For MRSA, values were 0.78% and 0.39% v/v, and for *E. coli*, 0.78% and 0.2% v/v, ranking oxaline as the second most potent DES alongside T-glycoline but displaying a slightly higher activity towards Gram-negative strain.

Overall, the obtained data follows the previously reported tendency of OA-DESs towards MRSA than to *E. coli* (Swebocki *et al*. 2024), but the observable effect (such as in a case of maloline and maline) is much more pronounced. This is interesting since the general mode of action of OA-DESs is through the destabilization of the pH equilibria, in the microenvironment(Hsiao and Siebert 1999) of the cells which likely leads to the denaturation of proteins(Lund, Tramonti and De Biase 2014), impairing the proper functioning of the cell. Such an approach is usually very effective and acts more like a “hammer” than a “scalpel” method, without major selectivity between the microbial species. However, as mentioned in the introduction, biofilms are extremely complex environments and are able to inhibit many chemical and physical factors on microbial cells. Since the differences between particular OA-DESs are much greater for biofilms, this partially confirms that the biofilm matrix is a key aspect limiting the effectiveness of OA-DESs and thus may alter the tendency of their antibacterial properties against biofilms when compared to the planktonic bacteria.

#### 3.1.2. Live/Dead assay (LD)

To complement MBEC assessment, we turned into the other experiment – that is, Live/Dead assay (LD). LD allows to image the living and dead bacteria using two fluorescent stains. Propidium iodide (PI) stains the dead cells red, while SYTO 9 stains both dead and live cells green. The superimposition of two images allows to roughly determine the cells survivability in the investigated environment.

After maturation and treatment with sub-MBEC99.9 OA-DESs’ concentrations, the biofilms were stained and imaged using fluorescence microscope. The images were then superimposed to obtain the final images of the biofilms.

As seen in the **Figure 3**, untreated matured biofilm of MRSA showed little-to-none of the fluorescence in the red channel, indicating just a small fraction of the dead cells. This is reflected in lower emission of the green channel as well, as the SYTO 9 stains both dead and live cells; with minor number of dead cells, the overall emission is also lower. The structure of the biofilm is slightly loose but definitely wrinkled with well-pronounced multilayered structure. The treatment of the biofilms with OA-DESs resulted in much stronger emission observed in the red channel, resulting in yellowish hue in the composed images. This is visible in all the images of OA-DES-treated biofilms, confirming the (partial) eradication of the biofilm-enveloped bacteria. The antibacterial properties of maline, which was the least potent antimicrobial, were also confirmed. This is well observed in scarce red emission and much higher emission of the green light when compared to other images. As shown in the previous experiment, maline, despite not being able to achieve MBEC was able to induce three-log-units decrease of the bacterial population, which explains the presence of the red emission. Interestingly, T-glycoline caused the appearance of the holes in the surface of the biofilm. Turning into *E. coli* biofilms, in the image presenting an untreated biofilm, we can clearly observe both red and green light emissions, indicating the presence of significant numbers of both dead as well as alive cells.

| MRSA |  | *E. coli* |  |
| --- | --- | --- | --- |
| control | glycoline | control | glycoline |

**Figure 3.**
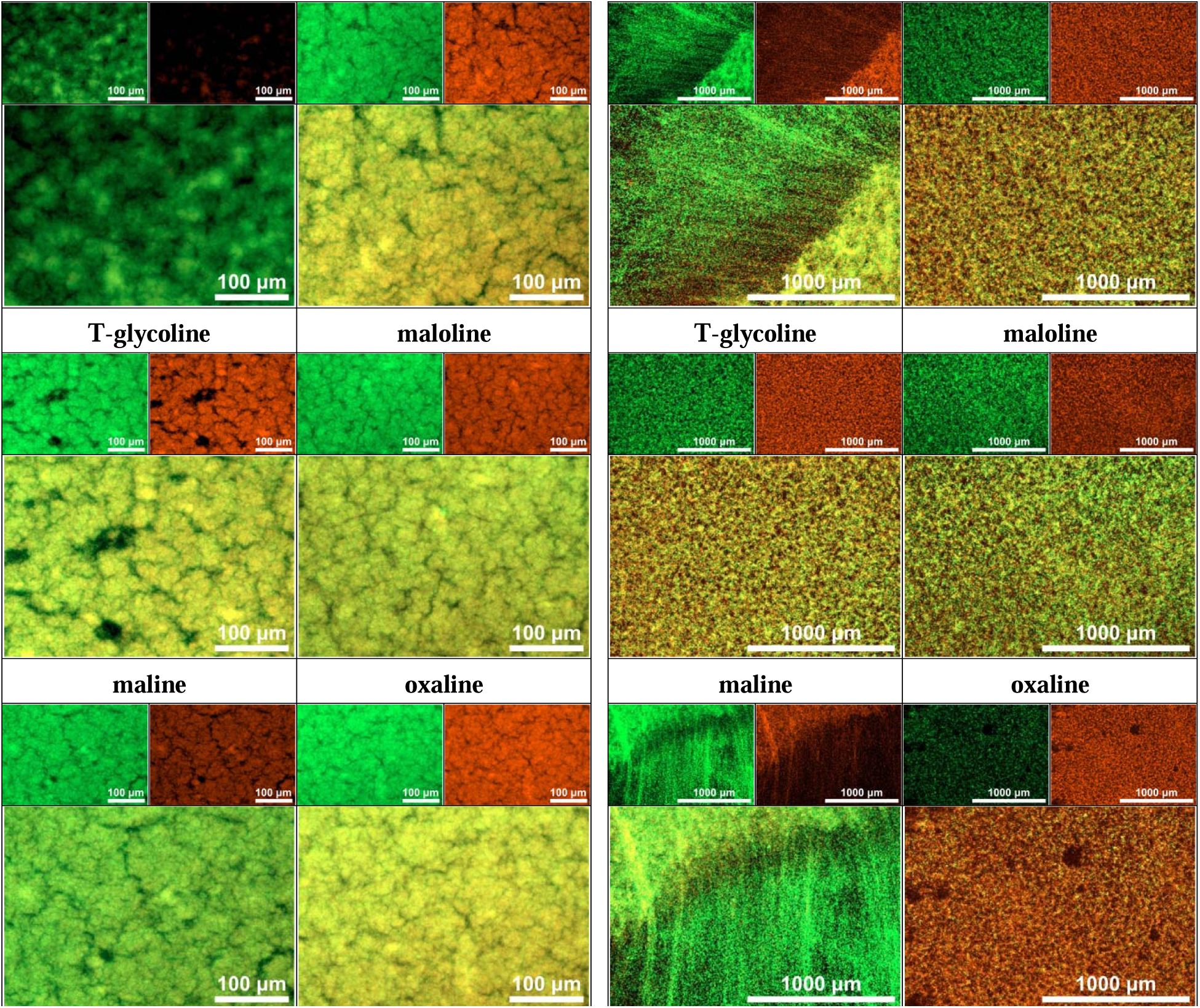
The Live/Dead assay of the MRSA (left) and *E. coli* (right) biofilms before and after the treatment. Small images indicate green and red channels, the bigger image is a superposition of both.

However, since the *E. coli* biofilm was maturing for 48 h, in contrast to 24 h for MRSA, the possible higher number of dead cells is expected and justified due to some cells already being in the late death phase. Interestingly, the morphology of *E. coli* biofilm displays less regularities when compared to MRSA, with visible thread-like strands of bacterial matter radially growing out of more condensed population of bacteria. After the treatment, two observations can be taken immediately. Firstly, not all the OA-DESs treatments had same outcomes. In the case of all the OA-DESs but maline, we observed a homogenization of the morphology of the biofilm – indicating structural changes of the biofilm caused by the DESs. Here, glycoline and T-glycoline displayed similar activity with significant number of dead cells visible in the images. Maloline was slightly less effective, presenting weaker emission of the red light. Oxaline, which was the most potent OA-DESs against *E. coli* also caused significant eradication of bacteria, as shown in the major emission of the red light, and reddish hue in the composed image. Another observation made is the presence of the non-treated biofilm morphology after treatment of the *E. coli* biofilm with maline – here, again the strand like structure is visible, radially propagating from the denser populations of bacteria. The above-mentioned results indicate that OA-DESs are effective eradicating agents, which can cause morphological changes to the biofilms as well. However, to draw direct conclusions about the nature of the interaction of OA-DESs with biofilm, additional investigation is advised – primary to verify the interaction of the EPS with OA-DESs as well as the pace of the eradication or the kinetics of the process.

### 3.2. Persister cells assay

As mentioned in the introduction, biofilms pose a direct danger to human-focused healthcare. However, many medical specialists and researchers are worried about subpopulation of the cells, which can effectively “hide” in the biofilms – that is, persister cells. Due to their ability to adapt to the inhibiting chemical factors, they can withstand the presence of the antibiotics in increased concentrations. Should we worry that this applies to OA-DESs as well? Based on the plausible mechanism, and harsh environment caused by the low pH of the OA-DESs (**Figure S3**), it may seem to not be the case. However, to truly confirm the effectiveness of OA-DESs, one has to verify emergence of persister cells in the treatment-like environment. To this end, planktonic bacteria were subjected to all the OA-DESs in their MBEC.

The treatment of MRSA (**Figure 5**) resulted in varied activity. Oxaline was the most effective OA-DESs with very high pace of the eradication of the bacteria, which occurred almost instantly. As for maloline, the population was stable for the first minutes, however, it slowly started to decrease, with more than 90% of decrease in the first 45 min. After this time, a decrease of the bacterial population began, which finally resulted in a total eradication after 2 h. Glycoline and T-glycoline displayed very similar patterns, with prolonged phase of stable level of the bacterial population reaching 24 h for glycoline, and 6 h for T-glycoline. In the case of the latter one, a total eradication also occurred after 24 h. Over the course of 48 h, no increase in the bacterial population was observed for all the OA-DESs, indicating lack of the emergence of the persister cells in the investigated time. Obviously, maline was rejected, as its MBEC against MRSA was not reached and further increase of its concentration would likely result in the major dilution of microbial medium, which could interfere with the results.

**Figure 5.**
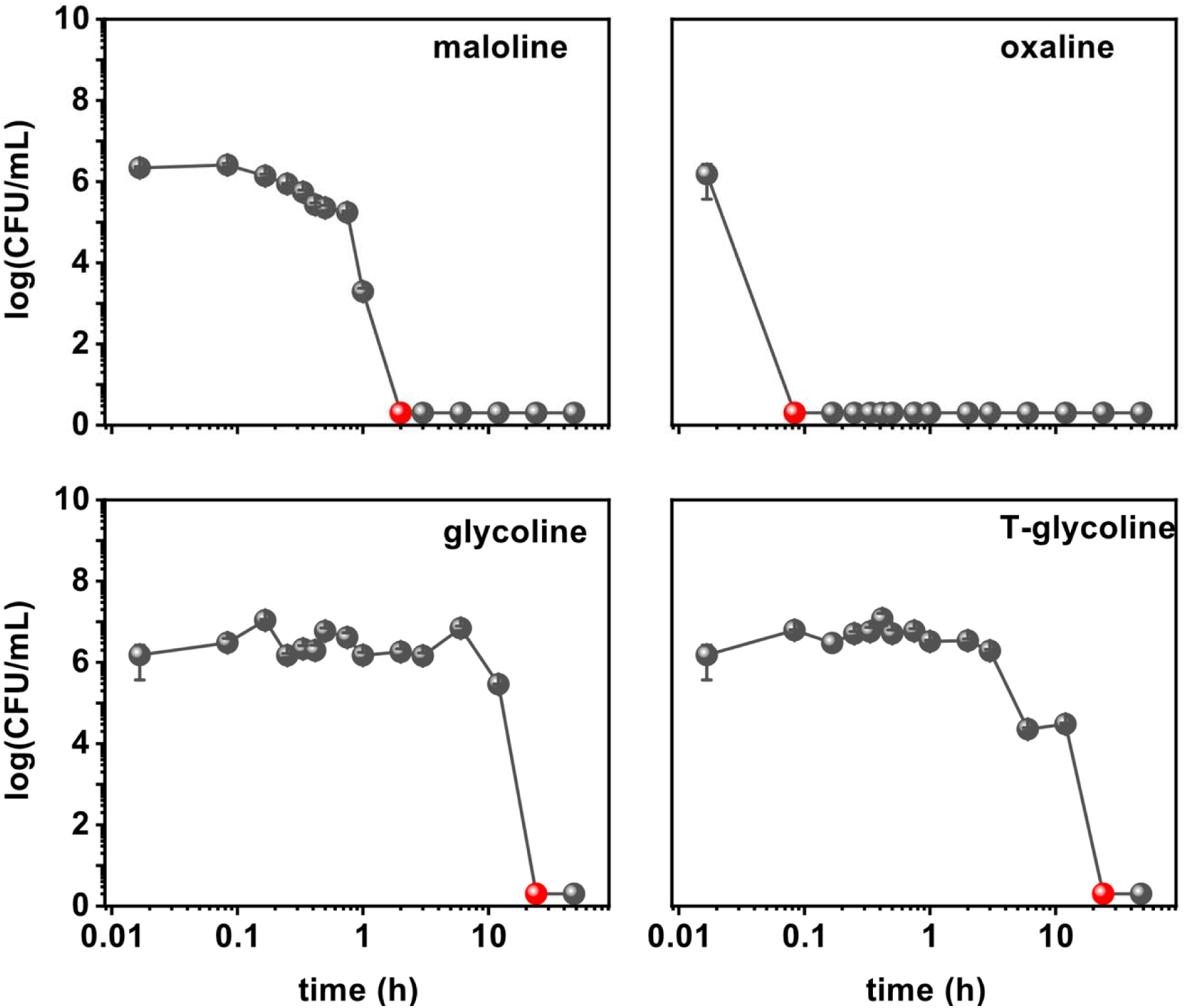
Persister cells assay for MRSA treated with MBEC of the investigated OA-DESs. Red points indicate a time-stamp when a total eradication of bacterial population occurred.

Much different results were observed for *E. coli* (**Figure 6**). Here, maline displayed the highest activity with almost immediate eradication of the bacterial cells, similarly to oxaline against MRSA. Glycoline, was the second OA-DES when it comes to the pace of the bacteria removal, with the time of eradication of 10 min., with more than a three-log-unit drop below 5 min. Maloline was also an effective antibacterial agent, with a time of 6 h needed to eradicate *E. coli* cells. Interestingly, shortly after the start of the experiment, the bacterial population decreased in number significantly with almost five-log-unit drop.

**Figure 6.**
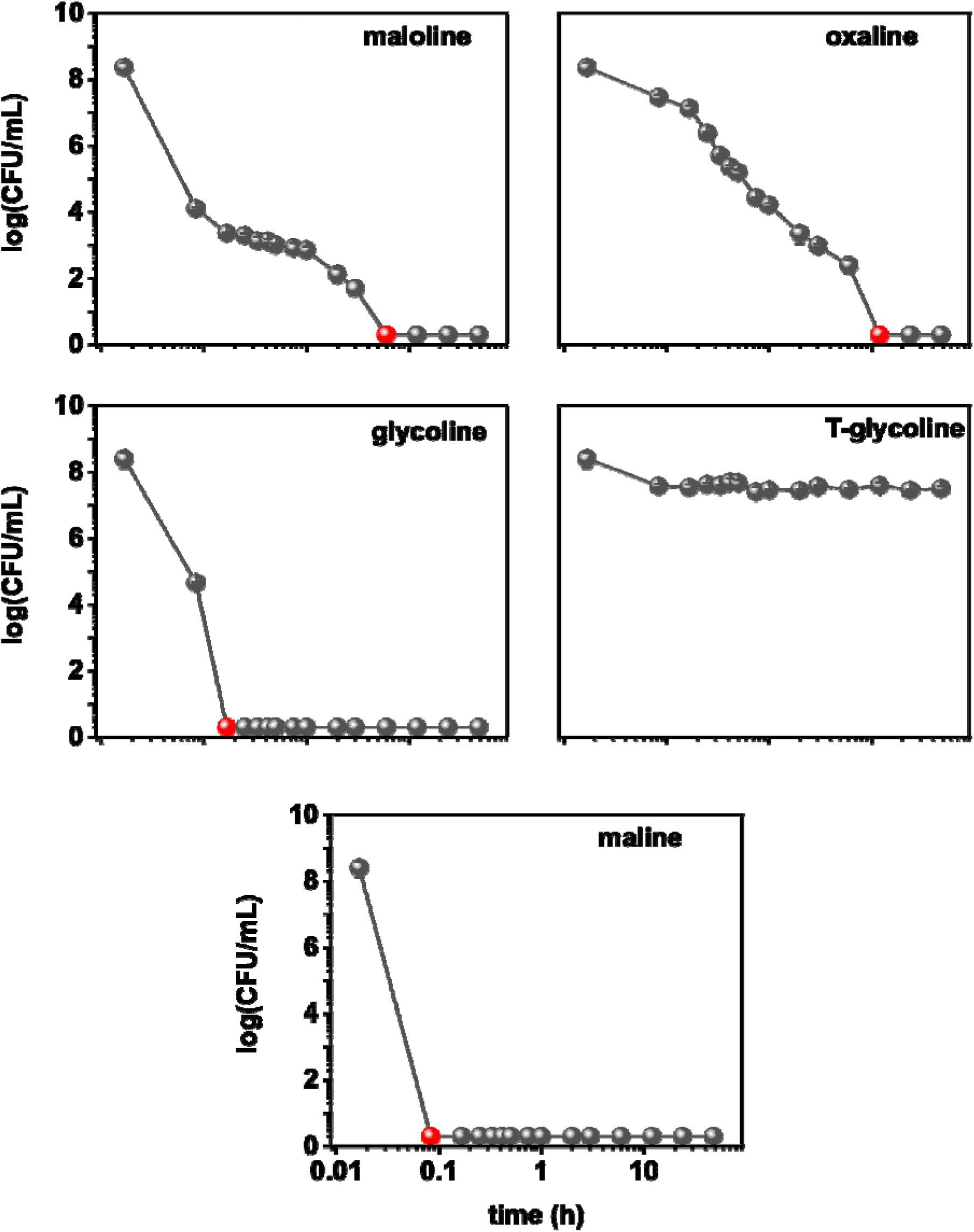
Persister cells assay for *E. coli* treated with MBEC of the investigated OA-DESs. Red points indicate a time-stamp when a total eradication of bacterial population occurred.

The number of the *E. coli* population was maintained over the course of almost 6 h, and finally dropped to zero, after that time. This may indicate that there exists a sub-population of *E. coli* that can withstand the increased concentration of maloline, although it does not display a resistance typical for persister cells. Oxaline, was the second least effective OA-DES. Over the course of 12 h, the population slowly decreased and only after that time reached zero – indicating total eradication. Such a behavior may indicate that oxaline has very slow kinetics. Worth noting is that our previous results of time-kill kinetics of oxaline against *E. coli* indicated otherwise(Swebocki *et al*. 2024), and show almost immediate action against *E. coli*. Though, as time-kill kinetics presented in the cited study assumes the use of another culture medium and much lower value of the initial cellular concentration for the test, the pace of eradication observed in this study is reasonable. Lastly, the use of T-glycoline, the last OA-DES tested, resulted in about one-log-unit decrease of the bacterial population. This level was maintained over the course of 48 h. As one would expect the cells to follow a standard growth curve (that is lag-phase, exponential phase, stationary phase, and death phase), the decrease of the bacterial population at the end of the experiment was not observed. This may indicate that bacteria slowed down their metabolism in the presence of the antimicrobial agent, which together with around 90% decrease of population confirms the presence of persister cells.

Overall, OA-DESs have been proved to be safe when it comes to the pontifical emergence of persister cells, which is crucial, in the aspect of biofilm eradication. In all but one case, OA-DESs did not cause the emergence of persister cells but varied in their pace of eradication which is also an important factor to be considered in future applications.

### 3.3. Inhibition of the fungal growth

#### 3.3.1. Yeast and mold control

Among different microbes, viruses and bacteria are often found responsible for posing high health-related risks. However, over recent years, many researchers are re-focusing their attention on other family of microbes – that is fungi. These microorganisms, similar to bacteria, are also very well-adapted to hospital environments – posing risk to immunocompromised patients. Owing to their metabolism and morphology, they are well-adapted to nutrient-limited and moist environment, which translates to their increased survivability in harsh environments.

With that in mind, we investigated the antifungal potency of the OA-DESs. To this end, we have selected three fungal strains: *Candida albicans*, *Candida auris* and *Aspergillus fumigatus*. *C. albicans* is a widely found inhabitant of the human microbiota;(Neville, d’Enfert and Bougnoux 2015) however, it can transition into a pathogenic state. Pathogenic *C. albicans* can cause a range of infections from mild mucosal to severe systemic diseases, especially in immunocompromised individuals(Ruhnke 2006; Mayer, Wilson and Hube 2013). Lately, great attention has been given to *C. auris*, which emerged as a multidrug-resistant strain. *C. auris* is capable of causing severe bloodstream-related infections, particularly in healthcare environments(Lockhart 2019). Lastly, *A. fumigatus* is a leading cause of invasive aspergillosis, primarily affecting individuals with compromised immune systems or pre-existing respiratory conditions(McCormick, Loeffler and Ebel 2010).

Different from bacteria for which a comparative study on the effect of the PSs and DESs formulations on their antibacterial effect was studied(Swebocki *et al*. 2024), no such report exists for the aforementioned fungal strains and OA-DESs investigated in this study. That being said, it is imperative to verify the antifungal activity of the OA-DESs in double parallel studies – that is, to compare the OA-DESs with non-OA-DESs (conventional DESs) as well as between their PSs and DESs themselves. Here, we have included four more DESs, which were ineffective in inhibiting bacteria(Swebocki *et al*. 2024) – reline (ChCl:urea 1:2), glyceline (ChCl:glycerol 1:2), ethaline (ChCl:ethylene glycol 1:2) and R-glyceline (L-arginine:glycerol 1:4).

The first investigated fungal strain was *C. albicans*. As seen in the **Figure S4**, most of the PSs had benign effect on the fungal cells, with only EG and TBACl displaying inhibiting effect. Apart from these two, OxA was the only component that caused the inhibition, but only at its highest concentration. When compared with DESs (**Figure 7**), we can make two interesting observations. Firstly, all of the DESs, except maloline, effectively inhibited the growth of the *C. albicans*. Most effective were T-glycoline and oxaline, which displayed remarkable activity at concentrations of 5% v/v. They were followed by maline at 5% v/v concentration which resulted in the fungal eradication. Apart from these three DESs, glycoline, ethaline, reline, and R-glyceline also effectively inhibited the growth of the *C. albicans*. However, another key observation is the significant increase of the OD_600_ in some cases. Such a phenomenon was displayed by glycoline (at 1% v/v), glyceline (5% v/v) maloline (whole range), R-glyceline (5% v/v) as well as maline (at 1% v/v). This indicates that these concentrations enhance the growth of the fungi – suggesting the proliferation effect.

**Figure 7.**
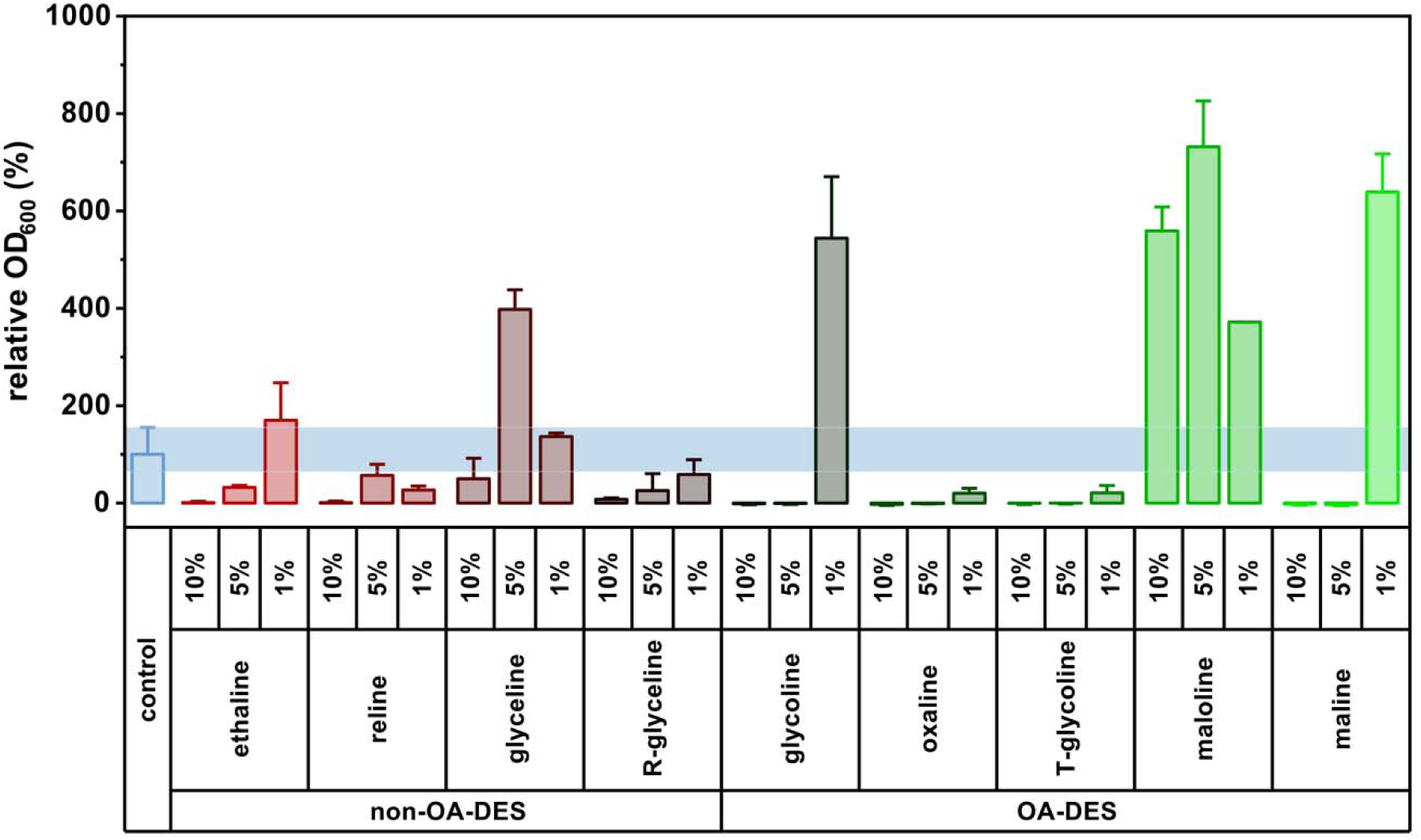
Antifungal assay of the investigated OA-DESs and non-OA-DESs against *C. albicans*.

Turning into microscopic images (**Figure S5**), certain morphological alterations were observed. Untreated *C. albicans* is characterized by its cluttered “cotton-like” filamentous form, as seen in the negative control – it is one of its virulence factors. Interestingly, in the case of OA-DESs, upon treatment with 1 % v/v, the clusters slightly increase in size (potential increase of the virulence factor), followed by the dispersion of the clusters into smaller aggregates. Similar behavior was observed for maloline, leading to the dispersion at 5 and 10% v/v, even though conversely to glycoline, never reached inhibition concentration. In the group of conventional DESs, reline (5% v/v) induced the growth of the fungal colonies, likely as a response to the environmental changes(Thompson, Carlisle and Kadosh 2011). However, further increase of the concentration led to significant inhibition. Ethaline displayed similar activity to glycoline – with 5% v/v concentration resulting in the (partial) dispersion of the filamentous colonies. Finally, glyceline also caused morphological changes of the colonies, firstly dispersing the cluttered colonies at 5% v/v, and further dispersion at 10% v/v, displaying the poorest overall activity in the group of conventional DESs.

In the case of *C. auris* (**Figure 8**), the antifungal activity was also observed in two groups of DESs. Notably, maline, oxaline, glycoline, and T-glycoline were very effective as the inhibition of the fungal population occurred at 5% v/v. Maloline was the only OA-DESs which did not display eradication at 10% v/v; however, a significant inhibition was still observed. R-glyceline and reline were the only non-OA-DESs that inhibited fungal growth at as low as 1% v/v. However, a significant inhibition was only spotted at the highest concentration. Ethaline followed the same pattern as reline, but with slightly higher OD_600_ values for lower concentrations. Glyceline was the only DESs that was not able to inhibit the fungal growth.

**Figure 8.**
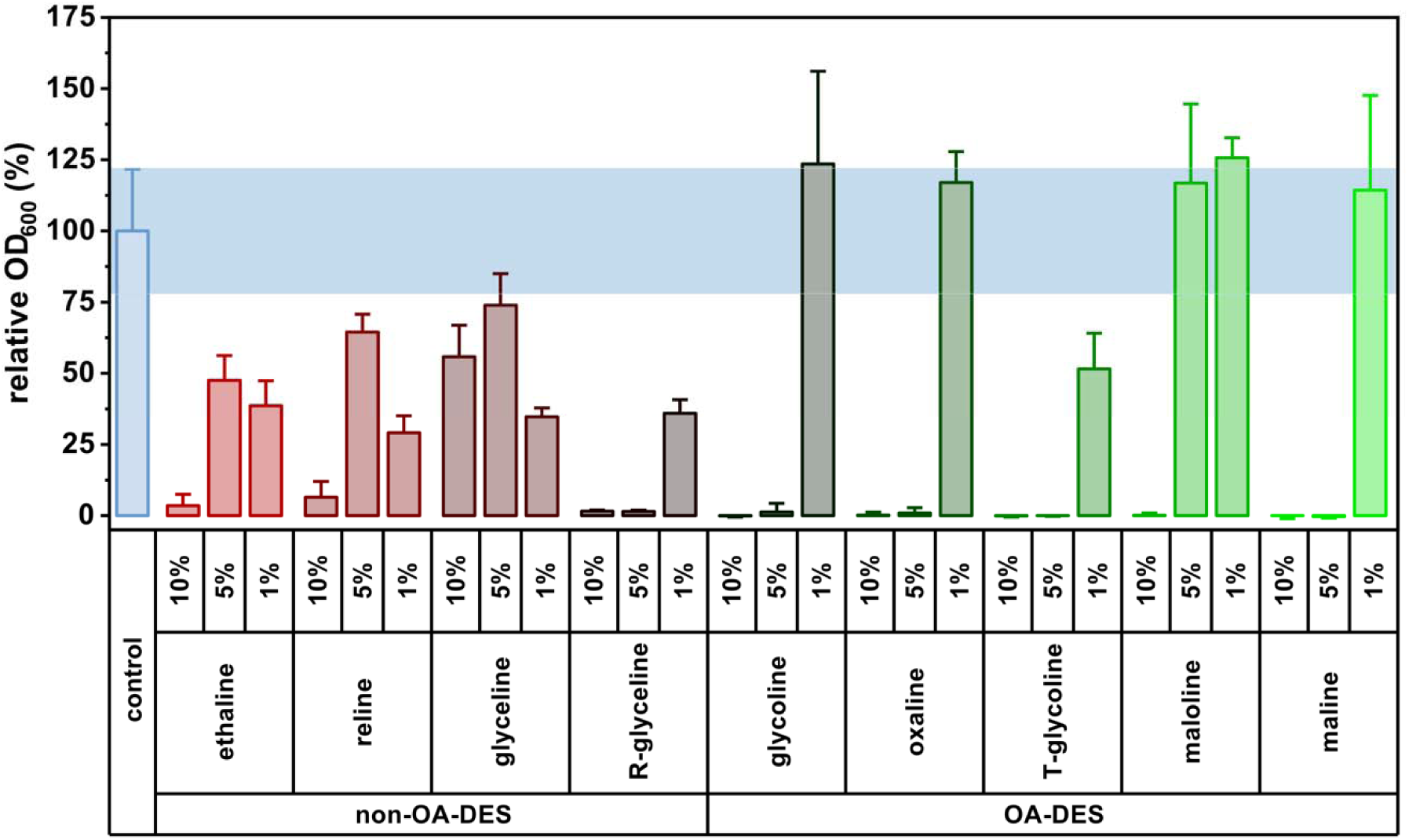
Antifungal assay of the investigated OA-DESs and non-OA-DESs against *C. auris*.

The antifungal assay of PSs (**Figure S6**) against *C. auris* is similar to the one for *C. albicans*. Most of the PSs showed little-to-none antifungal activity. However, only ethylene glycol (EG) and glycine (GLY) were able to subdue the fungal growth although no eradication was observed. Optical microscopy images (**Figure S7**) also showed morphological changes induced by DESs. A negative control revealed that *C. auris* maintained a well-dispersed form. Treatment with glycoline resulted in formation of small clusters of about 50-75 μm in size; however, their number is scarce suggesting significant inhibition of the fungus. Conversly, maloline induced filamentation to greater extent, with formation of clusters of around 150-200 μm in size – for both 5 and 10% v/v. For the non-OA-DESs, reline was the only DESs that caused some minor clusteration. The clusters are defined but seem less dense than for maloline. Similarly, ethaline caused minor filamentation, however most of clusters dispersed at higher concentration.

The final assessment covered *A. fumigatus* – a pathogenic mold. PSs screening showed that EG, GLY and OxA were the only substances able to cause partial inhibition (**Figure S8**), but not to the same extent as for both *Candida* strains. Though, when formulated into DESs they displayed significant antifungal activity (**Figure 9**). The most noticeable activity was recorded for maline, T-glycoline as well as glycoline, with all these OA-DESs effectively eradicating fungal strains at concentrations lower than 5% v/v. Maloline and oxaline, despite inhibiting the growth of *A. fumigatus*, were ineffective in a total eradication of the mold. Interestingly, in the group of non-OA-DESs, reline displayed remarkable activity, followed by ethaline. Glyceline and R-glyceline were the least effective DESs in this group, which reduced OD_600_ only by approx. 70%. Unlike for *Candida* strains, no morphological changes were induced to *A. fumigatus* (**Figure S9**).

**Figure 9.**
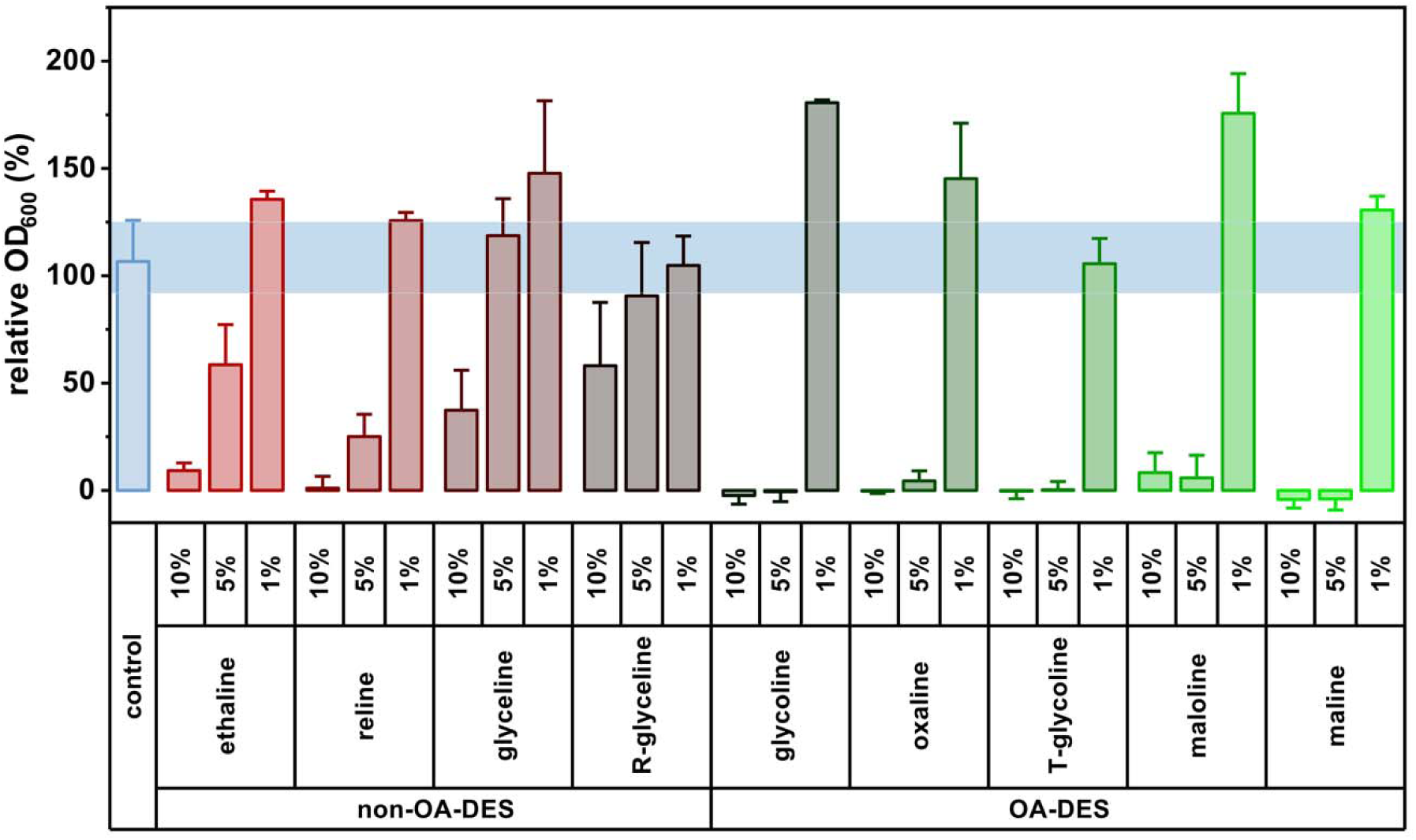
Antifungal assay of the investigated OA-DESs and non-OA-DESs against *A. fumigatus*.

It is worth noting that due to pH changes caused by OA-DESs (**Figure S3**), a standard colorimetric assay was impossible, as even the lowest concentration of DESs caused the change of color of resazurin, which is used in such types of tests. To avoid false-negative results, we opted for an adaptation of standard OD measurements. However, as fungi tend to cluster much more than bacteria, such measurements are burdened with much higher error (as visible by large error bars). That is why optical microscopy was implemented in parallel.

#### 3.3.2. Phytopathogenic fungi control

Apart from man-made environments, fungi are found in natural habits as well, posing threats to other organisms as well. For example, *V. inaequali*s is responsible for the most important disease in apple trees (apple scab)(Bowen *et al*. 2011). Similarly, *Z. tritici* is responsible for the most important disease in wheat (Septoria leaf blotch). For these two major crops, the diseases caused by these fungi lead to significant yield losses and require the use of fungicides. Different types of chemicals fungicides have been used for decades but have led to environmental pollution and cases of resistance.(Cordero-Limon *et al*. 2021; Hoffmeister *et al*. 2024) The fungal strains used were previously characterized for their resistance to fungicidal active substances used in agriculture. *V. inaequalis* Rs552 strain exhibited reduced sensitivity to triazoles fungicides family(Leconte *et al*. 2024) and *Z. tritici* T02596 was a multidrug resistant strain.

In the case of *V. inaequalis*, mycelium growth was not inhibited at the 0.1% concentration for all non-OA-DESs, while relatively weak inhibition was observed at the 1% concentration for R-glyceline and reline only (**Figure 10** and **S10**). For OA-DESs, mycelium growth inhibitions were observed from 0.1% concentration, with a slight inhibition for glycoline, T-glycoline, maloline and an important inhibition for oxaline (80%). For all OA-DESs, except maline, mycelium growth inhibition at 1% concentration was total (100%).

**Figure 10.**
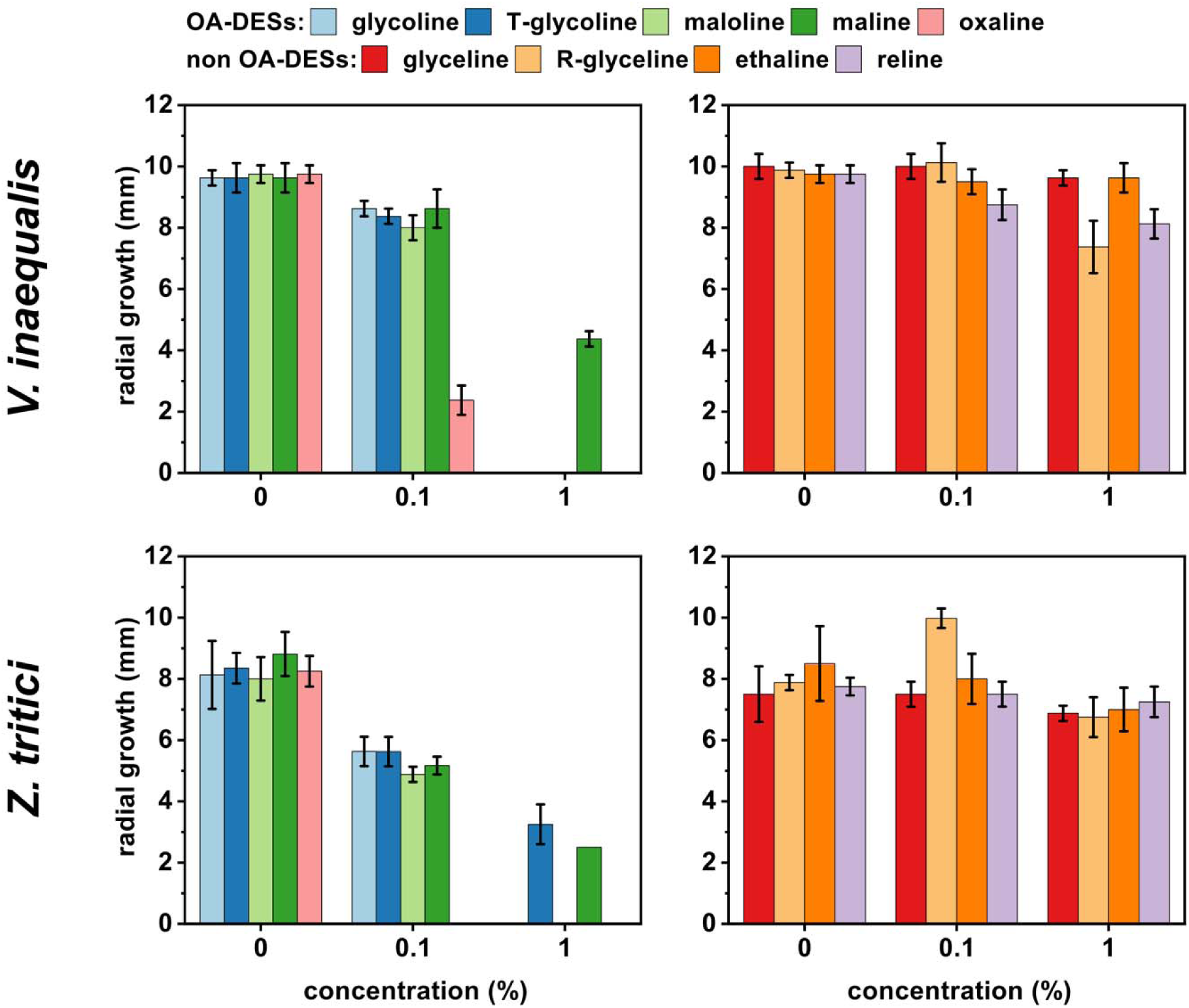
Antifungal assay of the investigated OA-DESs and non-OA-DESs against *V. inaequalis* (left) and *Z. tritici* (right).

In the case of *Z. tritici*, no mycelium growth inhibition was observed for all tested concentrations of non-OA-DESs (**Figure 10** and **S11**). For OA-DESs, mycelium growth inhibitions were observed from 0.1% concentration with total inhibition for oxaline only. For all OA-DESs, except T-glycoline and maline, mycelium growth inhibition at 1% concentration was total (100%).

For both fungi, we generally observed a lack of inhibition of mycelial growth for non OA-DESs which are neutral or basic pH compounds (**Figure S3**), whereas a stronger inhibition was observed for OA-DESs which are acidic compounds. For example, oxaline was showed strong inhibition at 0.1% concentration. The pH of oxaline at this concentration is 3 (pH = 1 at the 1% concentration). It seemed possible the acidification of the culture medium caused by oxaline was one of the reasons for its antifungal activity, especially as there was no total inhibition of maline at 1% (maline being one of the least acidic of the OA-DES compounds).

Previous works were already demonstrated antifungal activities of natural substances, as an alternative to chemical fungicides, against the used *V. inaequali*s and *Z. tritici* strains. However, this is the first time that antifungal activities have been shown with OA-DES compounds against these pathogens.

## 4. Conclusion

This study explored the antimicrobial efficacy of organic acid-based deep eutectic solvents (OA-DESs) against biofilms, persister cells, and fungi. The minimum biofilm eradication concentration (MBEC) assays showed that most OA-DESs eradicated MRSA and *Escherichia coli* biofilms at concentrations below 1% v/v. Notably, T-glycoline achieved complete eradication at just 0.39% v/v for both of the strains, while oxaline showed MBEC values as low as 0.78% v/v for MRSA and 0.2% v/v for *E. coli*. The Live/Dead assay further confirmed biofilm eradication, with extensive bacterial cell death and significant morphological changes in both strains. Importantly, all OA-DESs except one did not induce persister cell formation, with total eradication times ranging from 10 minutes to 24 hours, depending on the DES and bacterial strain. Beyond bacteria, OA-DESs displayed potent antifungal activity. Against *Candida albicans* and *Candida auris*, T-glycoline and oxaline were most effective, eradicating fungal populations at just 5% v/v, while maloline was the least effective and even promoted the growth of *C. albicans*. Similarly, against *Aspergillus fumigatus*, maline, glycoline, and T-glycoline showed complete eradication at <5% v/v. In phytopathogenic fungi, *Venturia inaequalis* and *Zymoseptoria tritici* exhibited 80-100% inhibition at 1% v/v for most OA-DESs, with oxaline achieving total growth suppression even at 0.1% v/v. Overall, OA-DESs provide a highly effective, broad-spectrum antimicrobial approach with potential applications in healthcare, industry, and agriculture. Their ability to eradicate biofilms, prevent persister cell formation, and inhibit fungal growth at low concentrations underscores their potential as next-generation antimicrobial agents. Future research should focus on refining their mechanisms of action and optimizing their real-world applicability.

## Financing and acknowledgments

T.S. would like to Gdańsk Tech for financing within Nobelium grant from IDUB funds (5/1/2024/IDUB/I.1a/No). Part of this research was done during the doctorate of T.S. which was financed by the former French Ministry of Higher Education, Research, and Innovation (MESRI). The CNRS, the University of Lille, and the Hauts-de-France region are acknowledged for financial support.

## Conflict of interest

The authors do not declare any conflict of interest

## Data availability statement

The data obtained in this research were deposited in the Institut d’Electronique de Microélectronique et de Nanotechnologie and are readily available by request.

## Author contributions

conceptualization, T.S., R.B.; data curation, T.S., K.C., C.B., J.M., J.J.; formal analysis, T.S.; funding acquisition, K.H., R.B.; investigation, T.S., K.C., C.B., J.M.; project administration, T.S., R.B., M.P.; resources, A.S., B.S., R.B., M.P.; supervision, R.B.; validation, T.S., C.B., J.M; visualization, T.S.; writing – original draft, T.S, A.M.K., J.M., A.S.; writing – review and editing, T.S., A.M.K., K.C., J.M., M.S.M., J.J., A.S., B.S., R.B., and M.P.

All authors read and agreed to the published version of the manuscript.

## SUPPORTING INFORMATION

**Figure S1.**
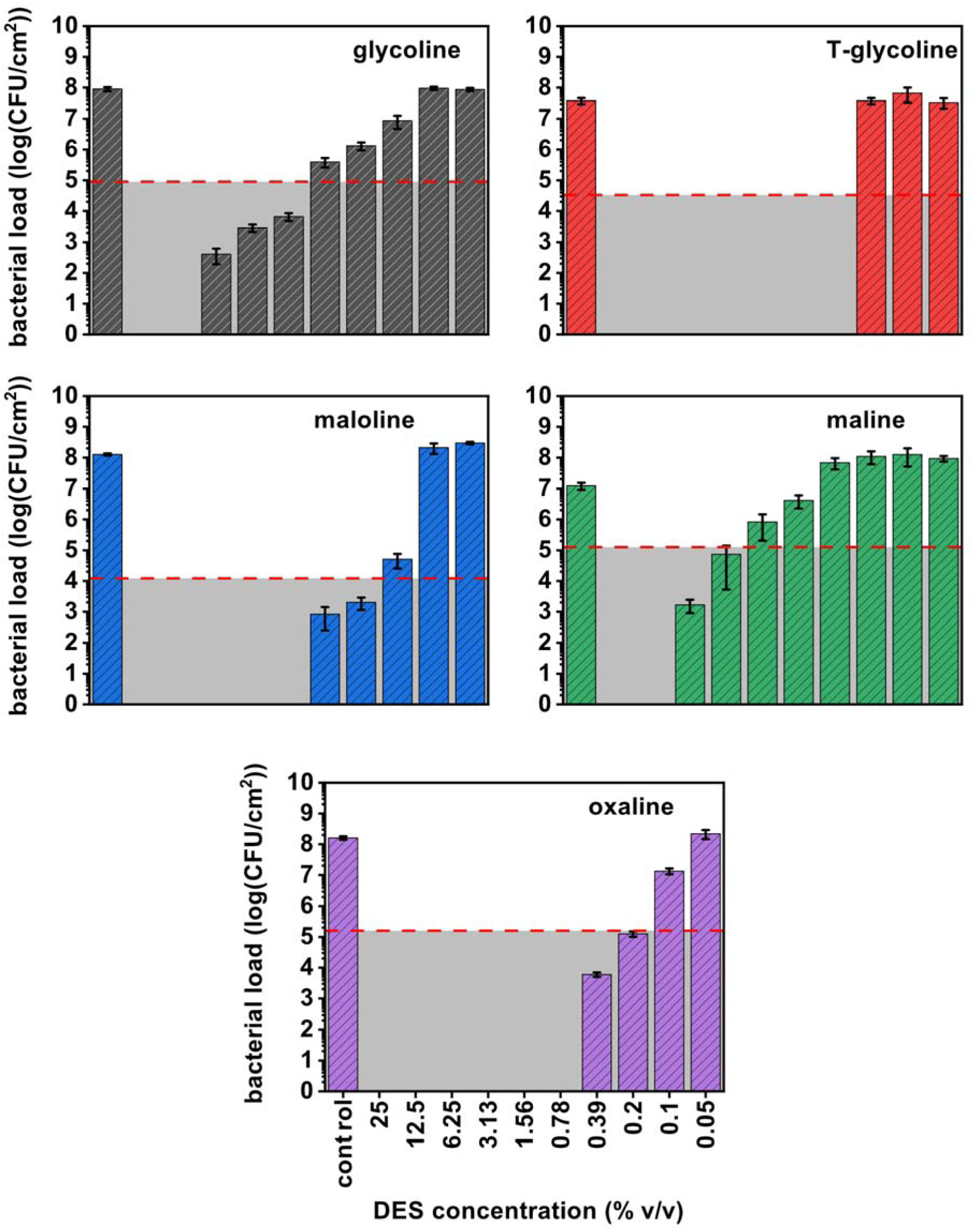
MBEC assay of the investigated OA-DESs against *E. coli*. Red dashed line indicates bacterial load corresponding to three-log-unit drop (MBEC99.9).

**Figure S2.**
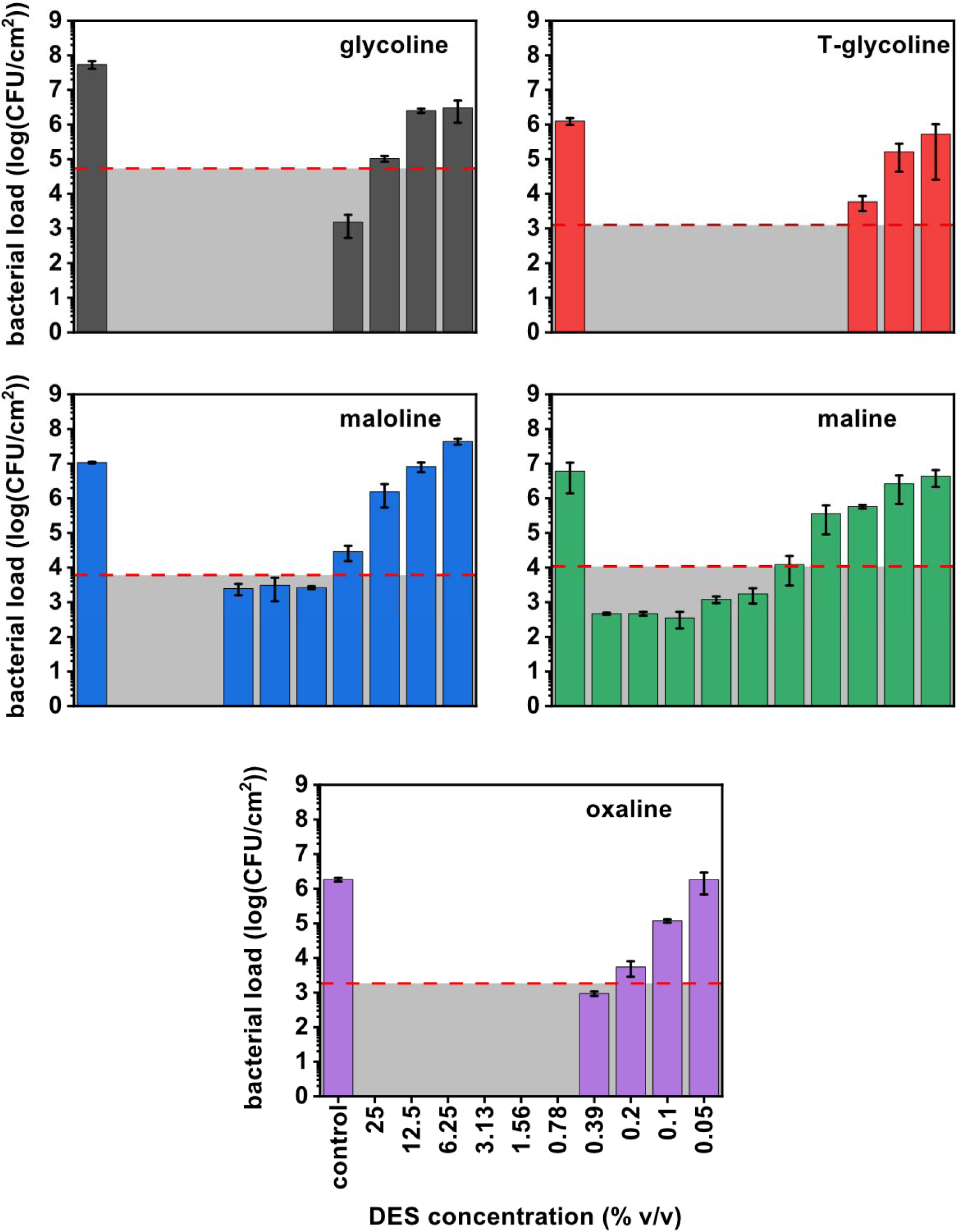
MBEC assay of the investigated OA-DESs against MRSA. Red dashed line indicates bacterial load corresponding to three-log-unit drop (MBEC99.9).

**Figure S3.**
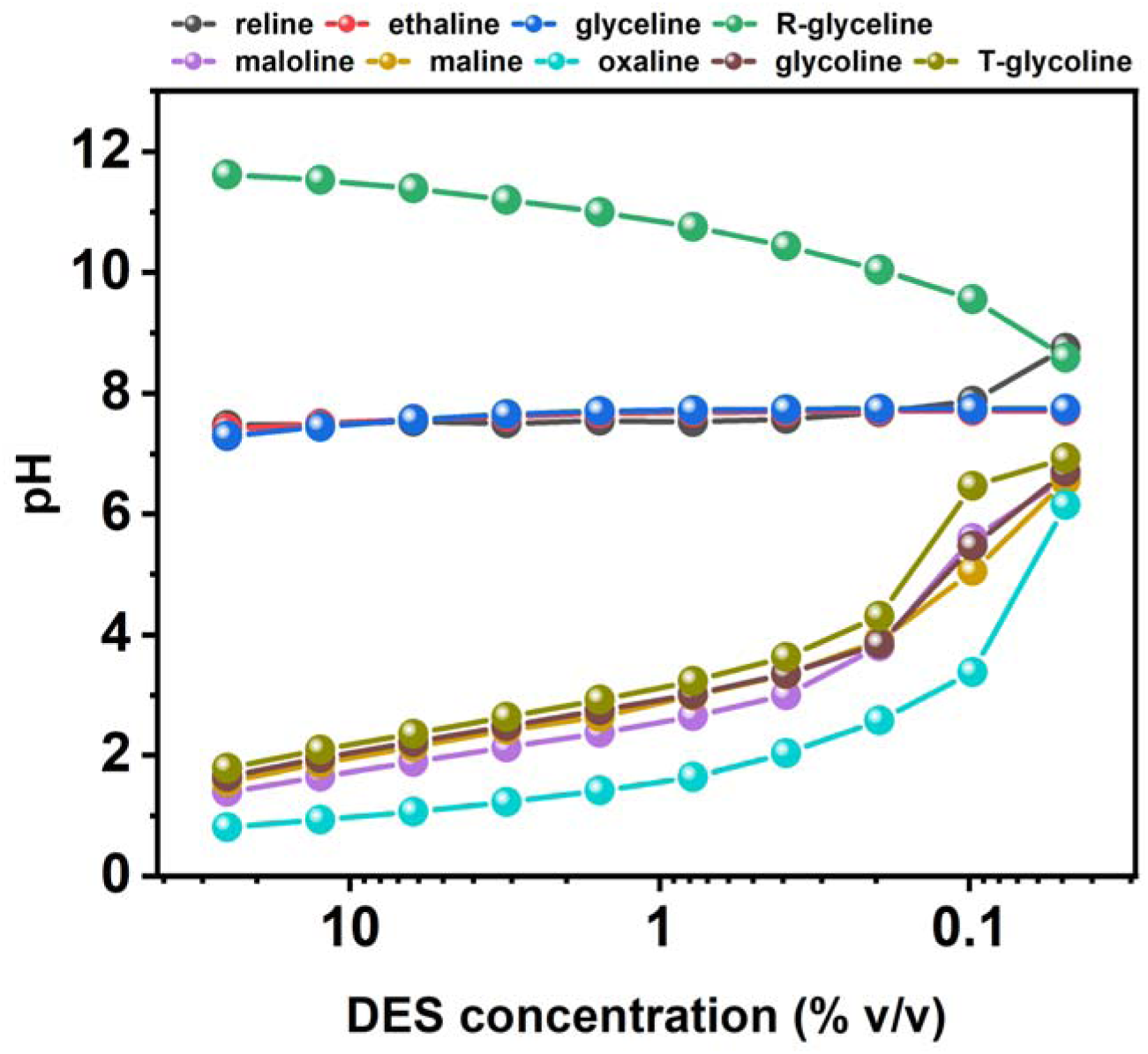
pH of the DES/water systems in the function of DESs’ concentration.

**Figure S4.**
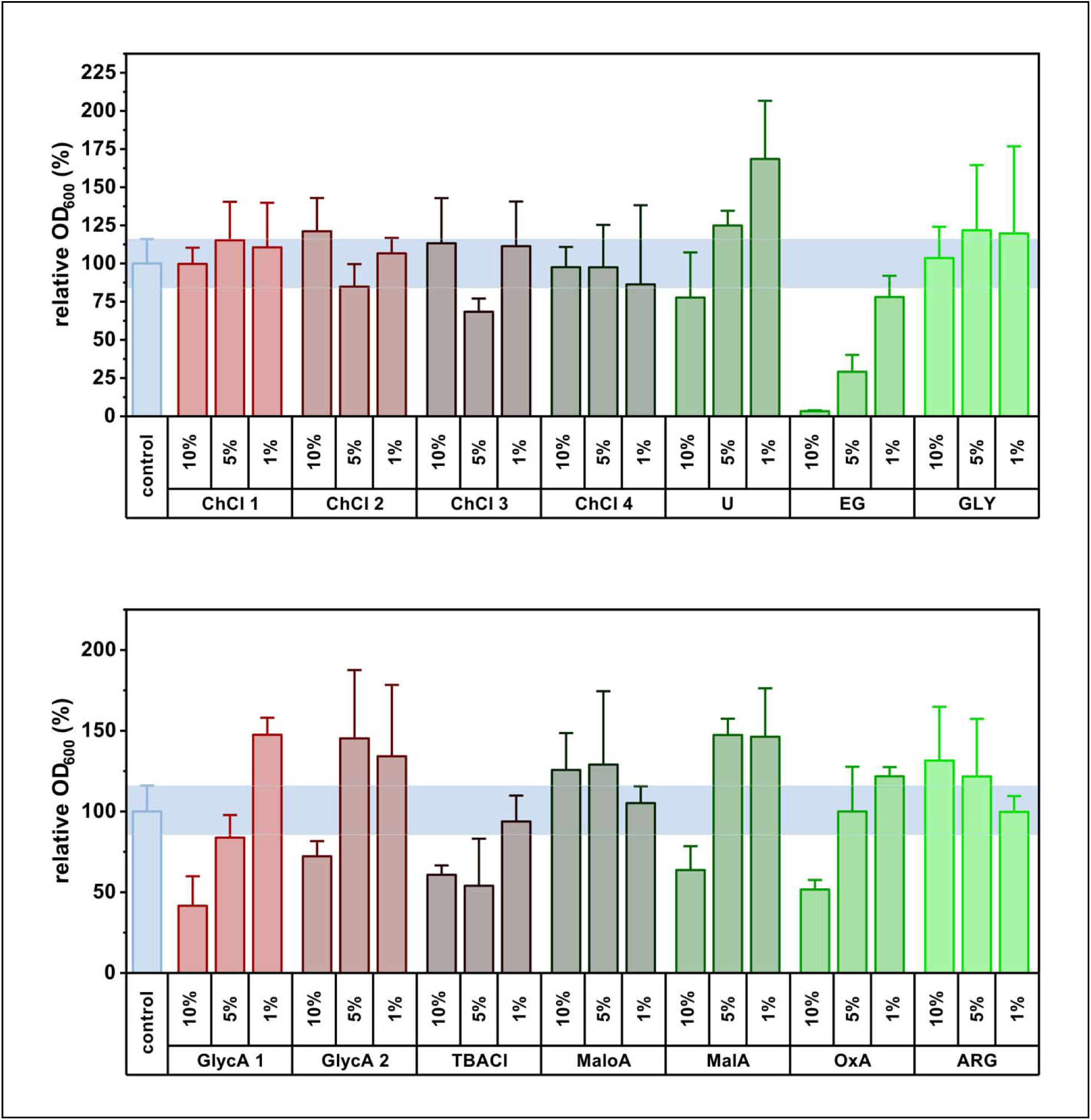
Antifungal assay of the PSs of the investigated DESs against *C. albicans* in 1, 5, and 10 % v/v equivalents (“ChCl #” indicates the concentration of ChCl corresponding to that of # = 1 – glycoline, maline and ethaline 2 – maloline and reline, 3 – glyceline, 4 – oxaline; “GlycA #” indicates the concertation of GlycA corresponding to that of # = 1 – glycoline, 2 –T-glycoline).

**Figure S5.**
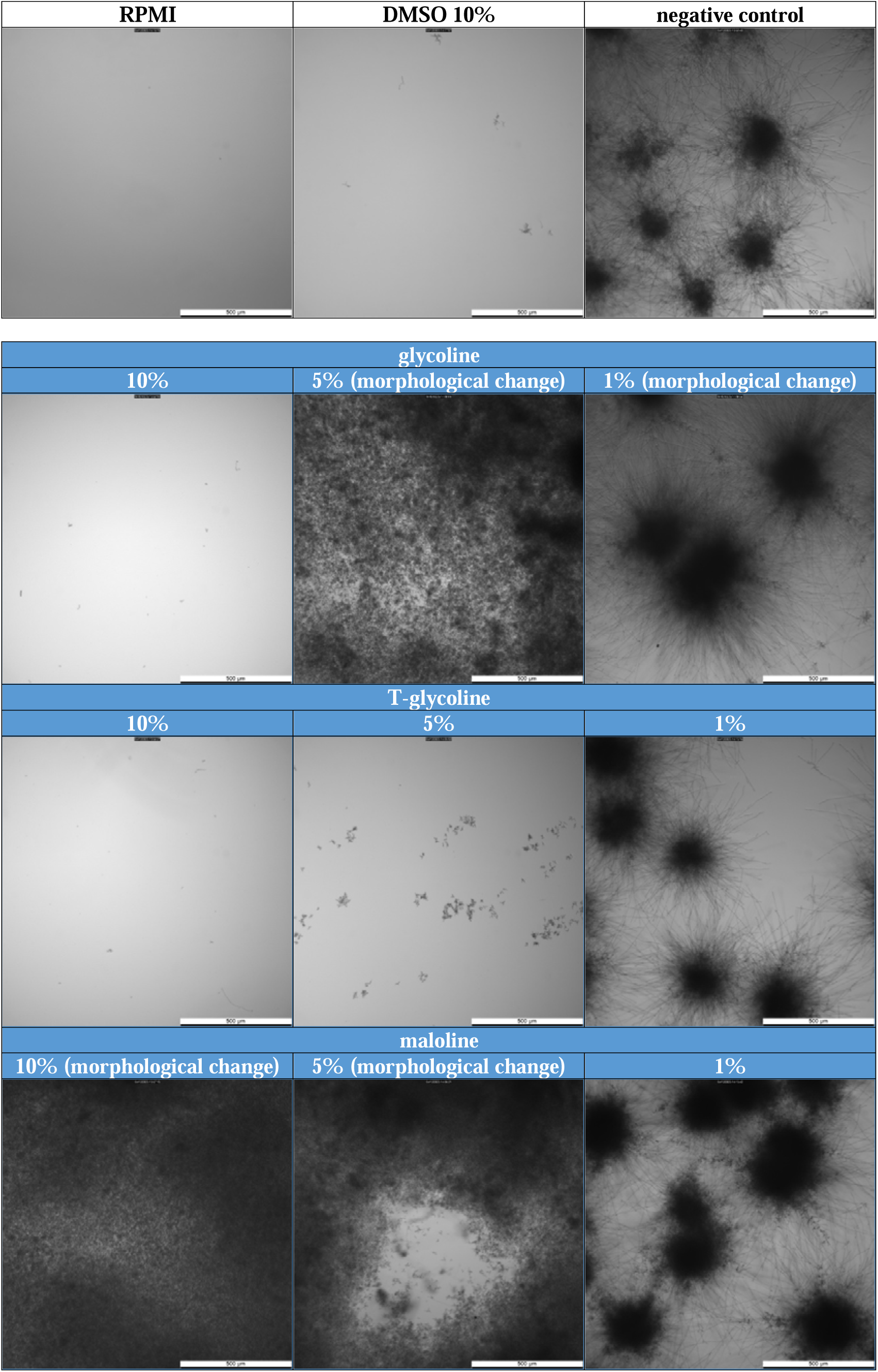
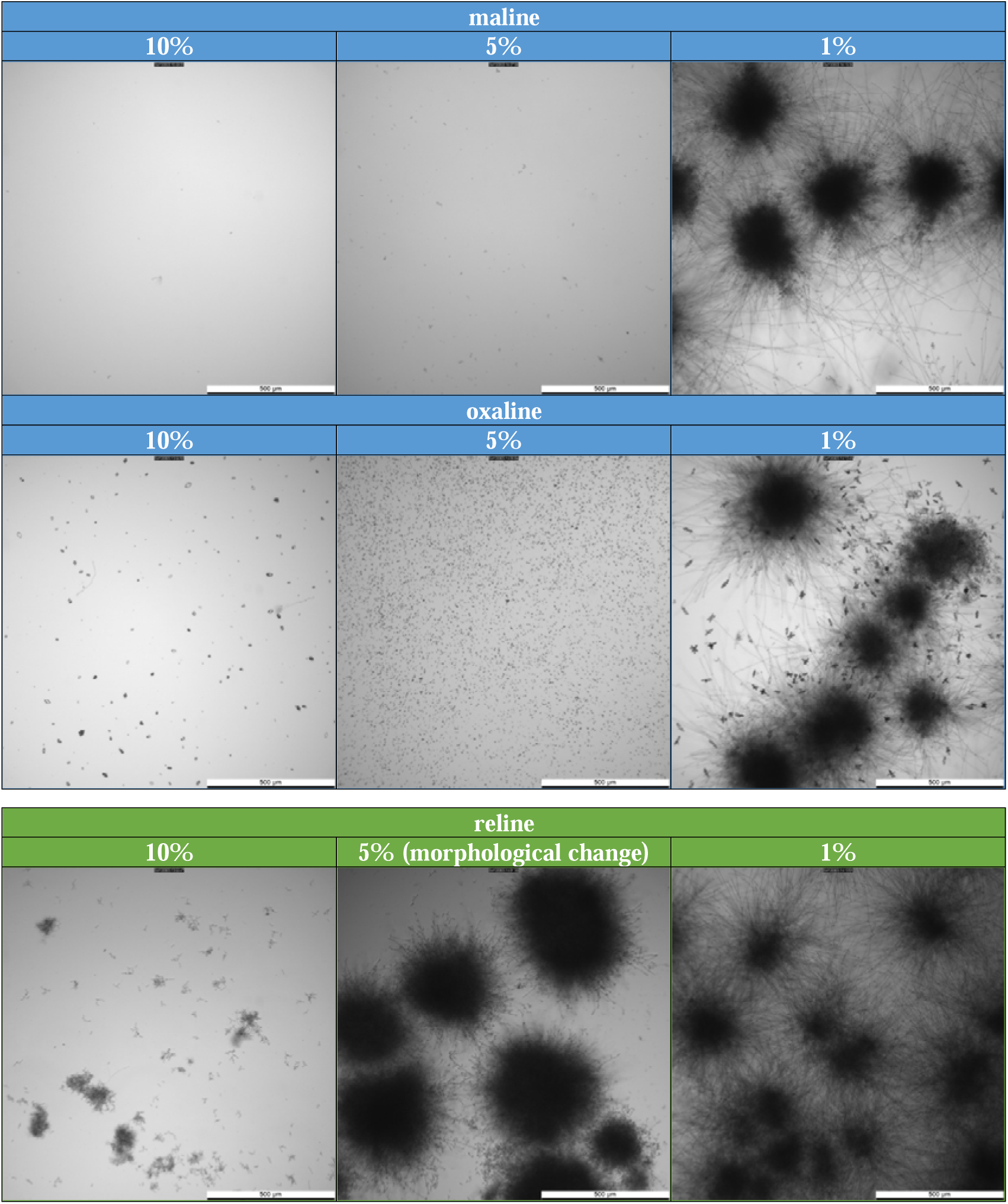
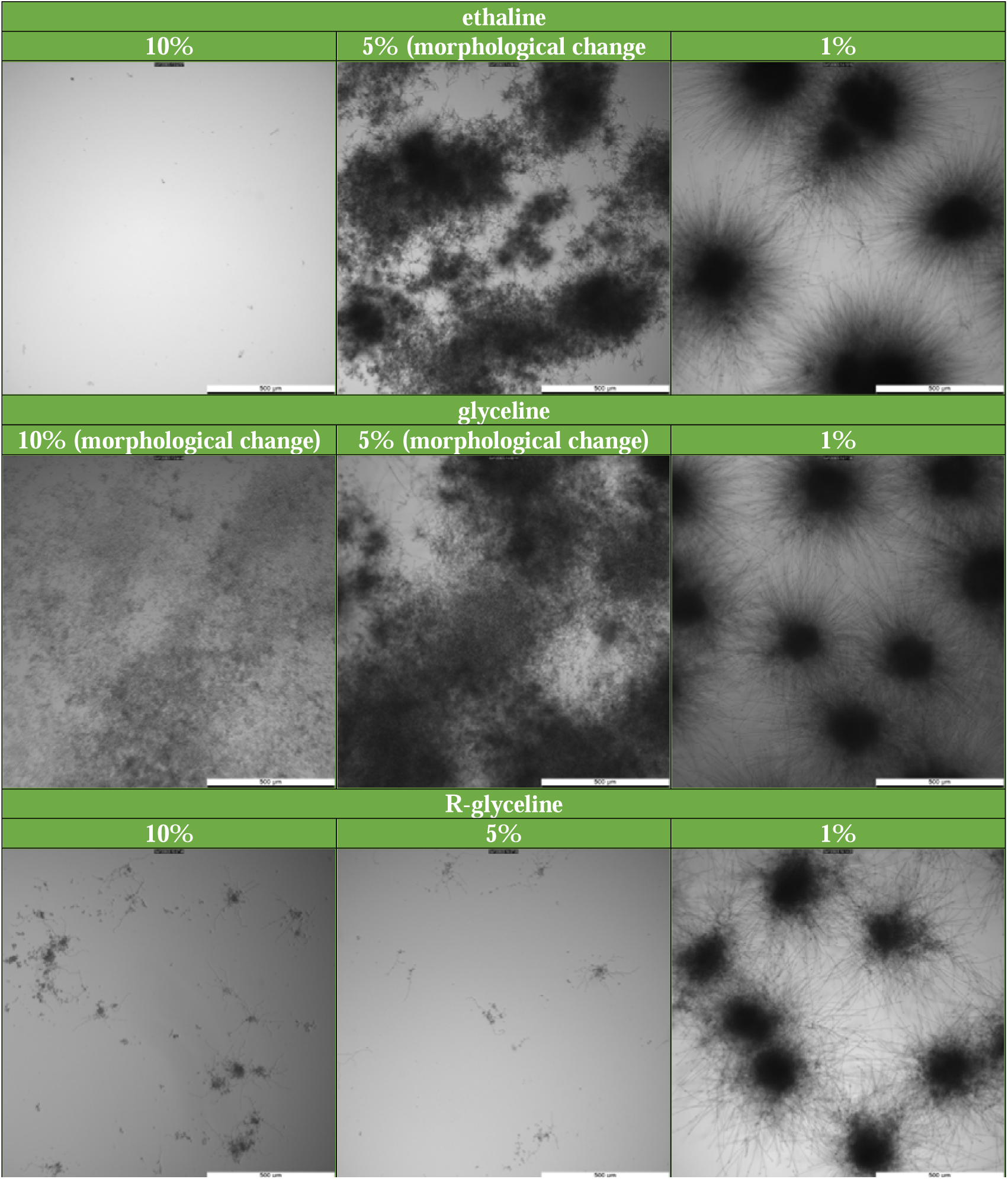
Optical microscopy images of *C. albicans* in the presence of the investigated DESs.

**Figure S6.**
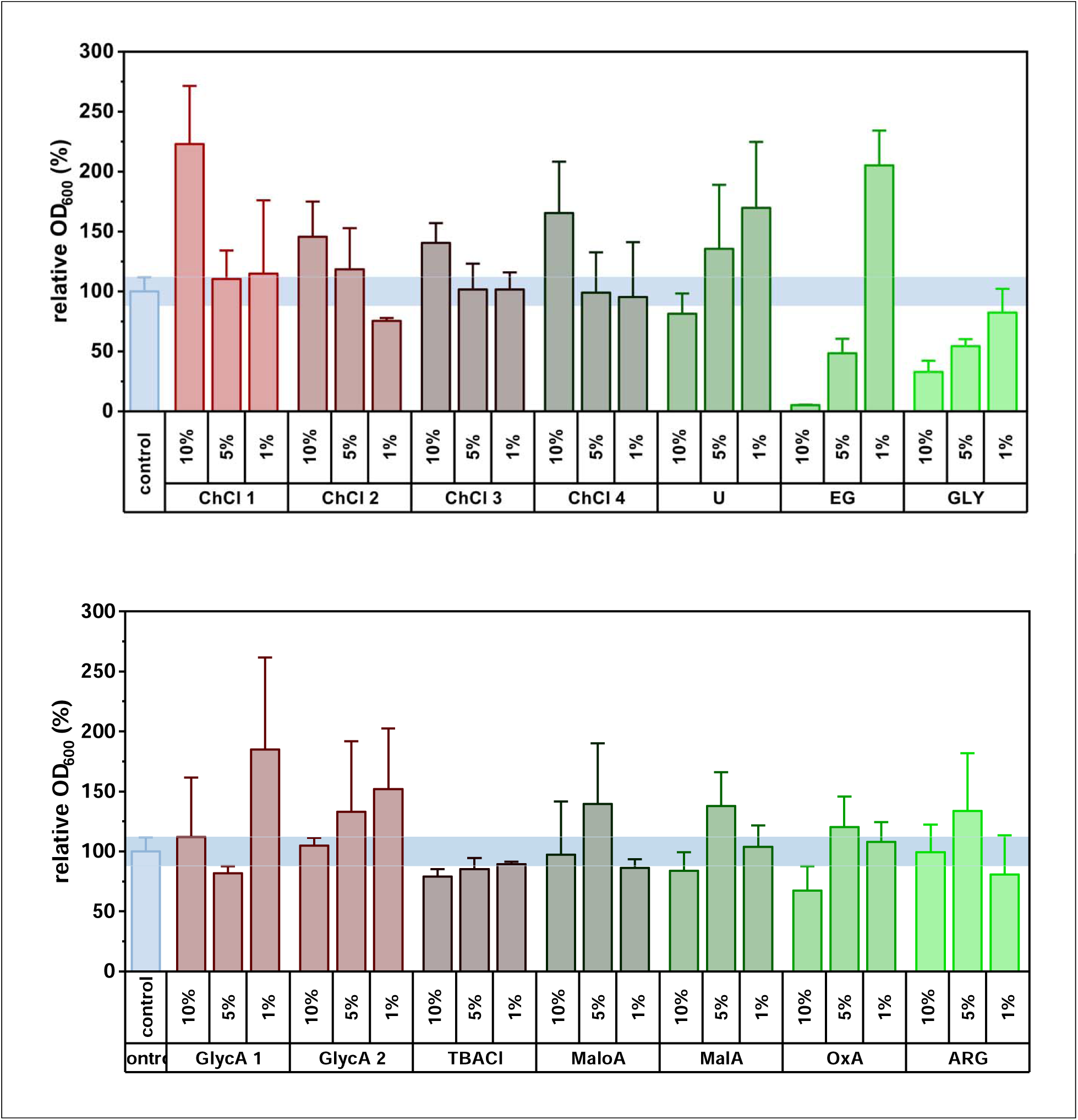
Antifungal assay of the PSs of the investigated DESs against *C. auris* in 1, 5, and 10 % v/v equivalents (“ChCl #” indicates the concentration of ChCl corresponding to that of # = 1 – glycoline, maline and ethaline 2 – maloline and reline, 3 – glyceline, 4 – oxaline; “GlycA #” indicates the concertation of GlycA corresponding to that of # = 1 – glycoline, 2 – T-glycoline).

**Figure S7.**
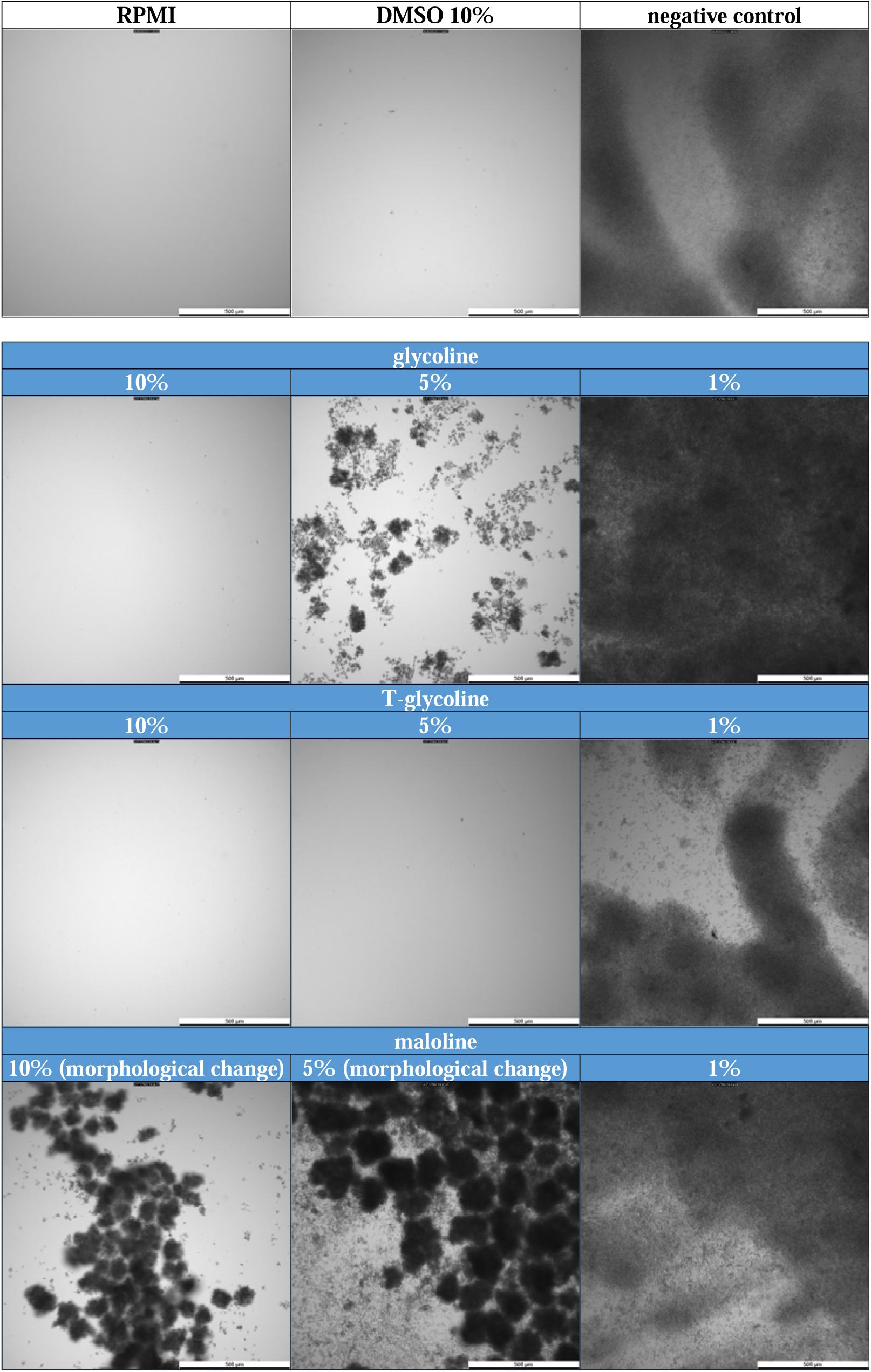
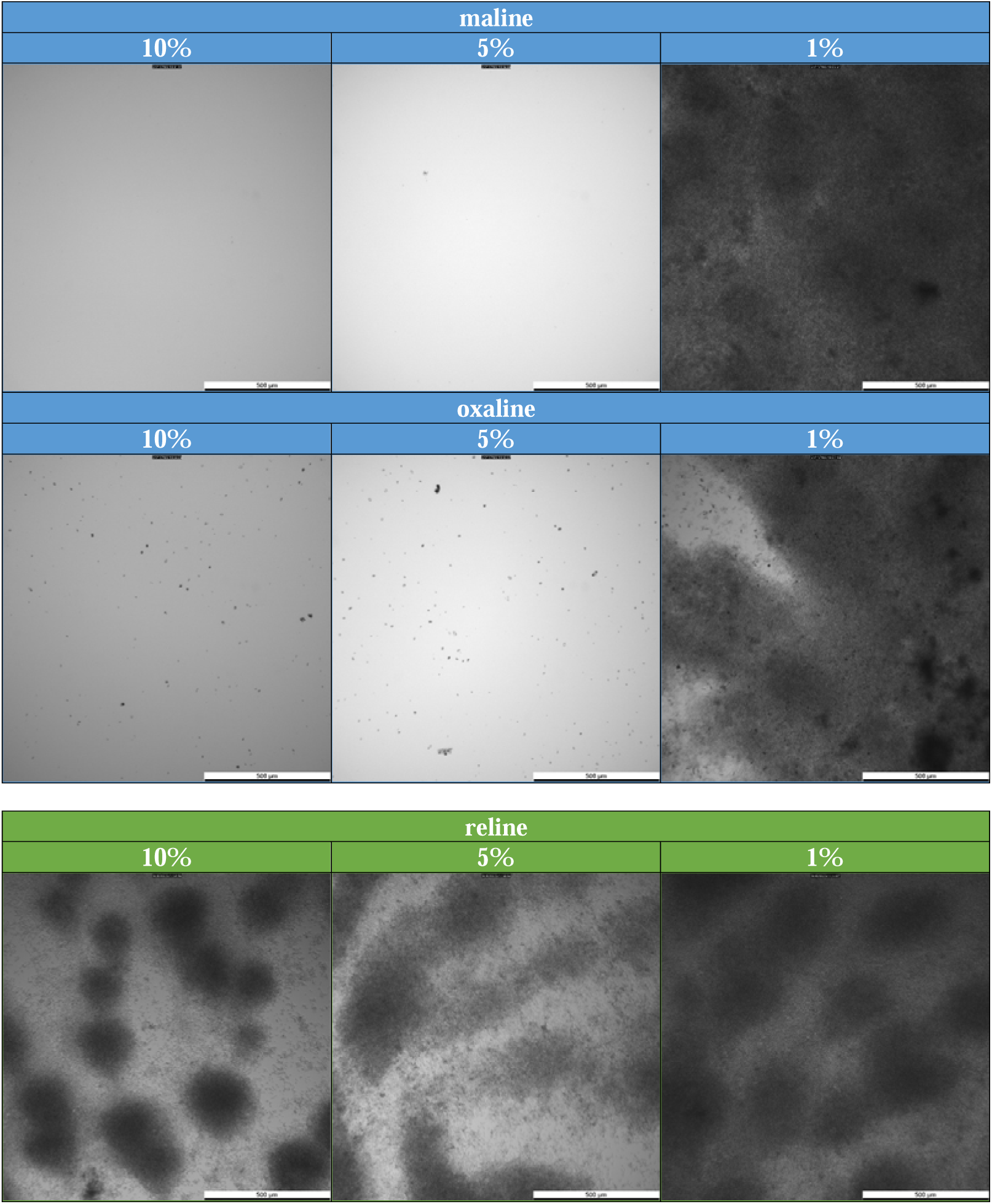
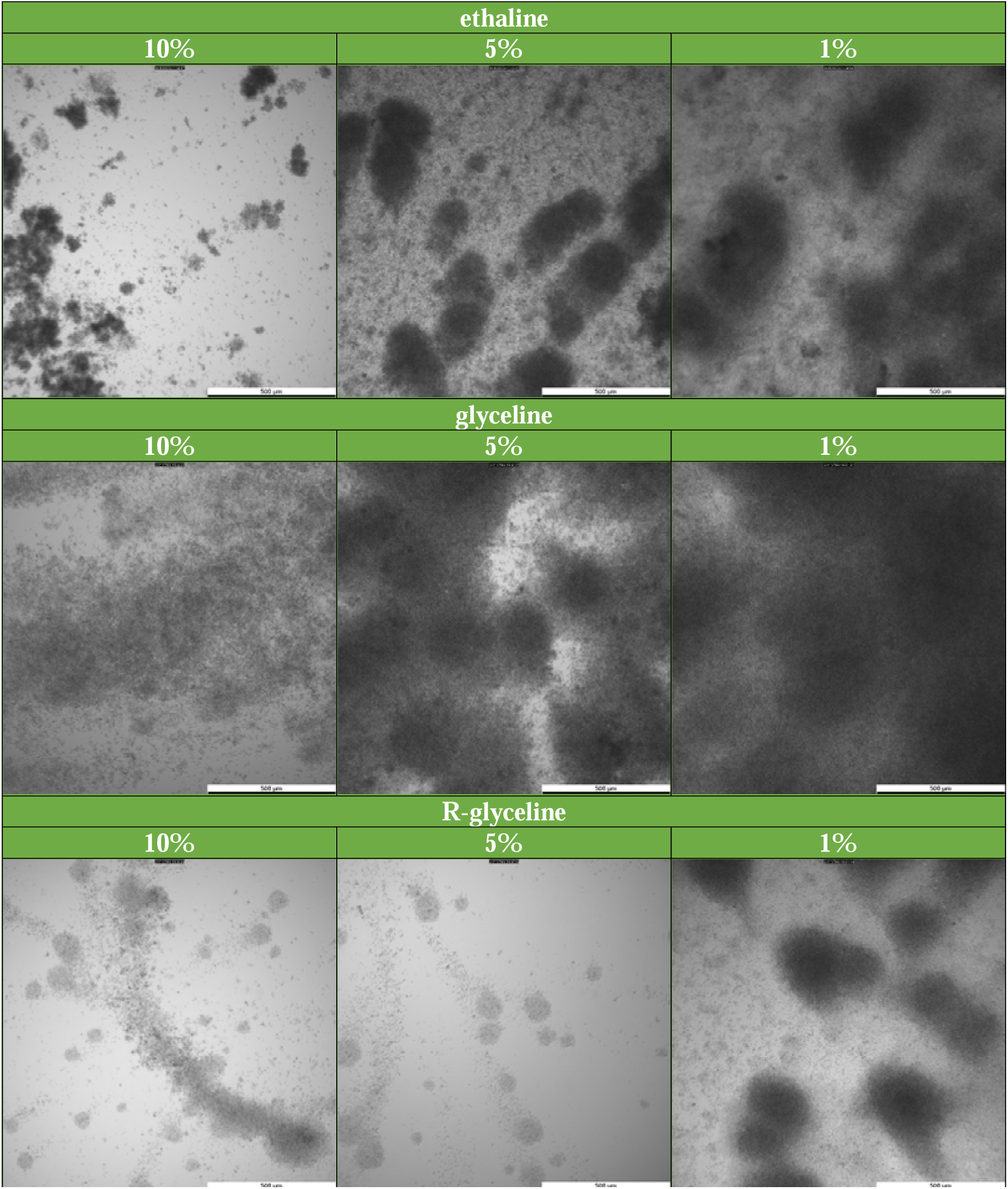
Optical microscopy images of *C. auris* in the presence of the investigated DESs.

**Figure S8.**
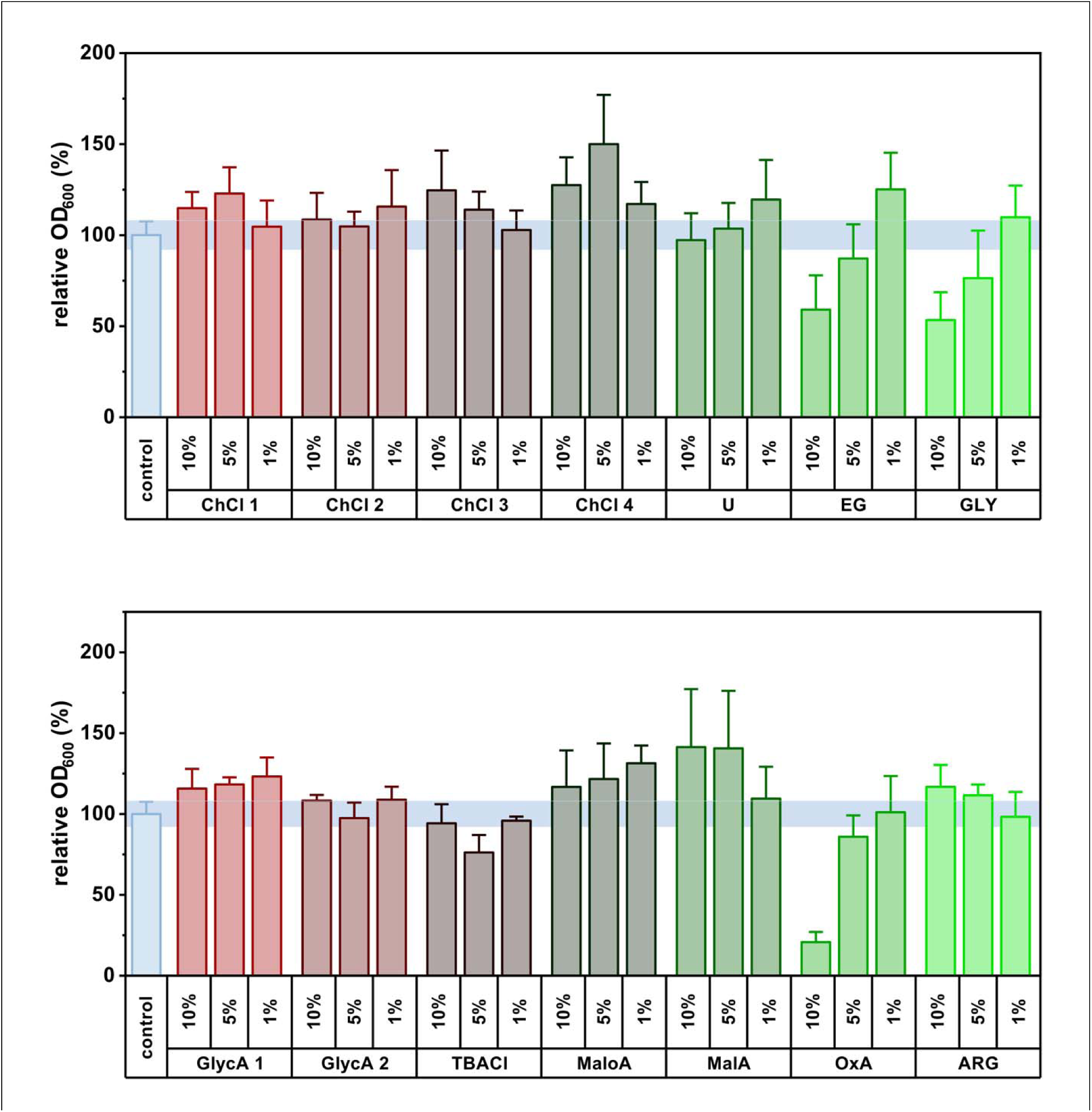
Antifungal assay of the PSs of the investigated DESs against *A. fumigatus* in 1,5, and 10 % v/v equivalents (“ChCl #” indicates the concentration of ChCl corresponding to that of # = 1 – glycoline, maline and ethaline 2 – maloline and reline, 3 – glyceline, 4 – oxaline; “GlycA #” indicates the concertation of GlycA corresponding to that of # = 1 – glycoline, 2 –T-glycoline).

**Figure S9.**
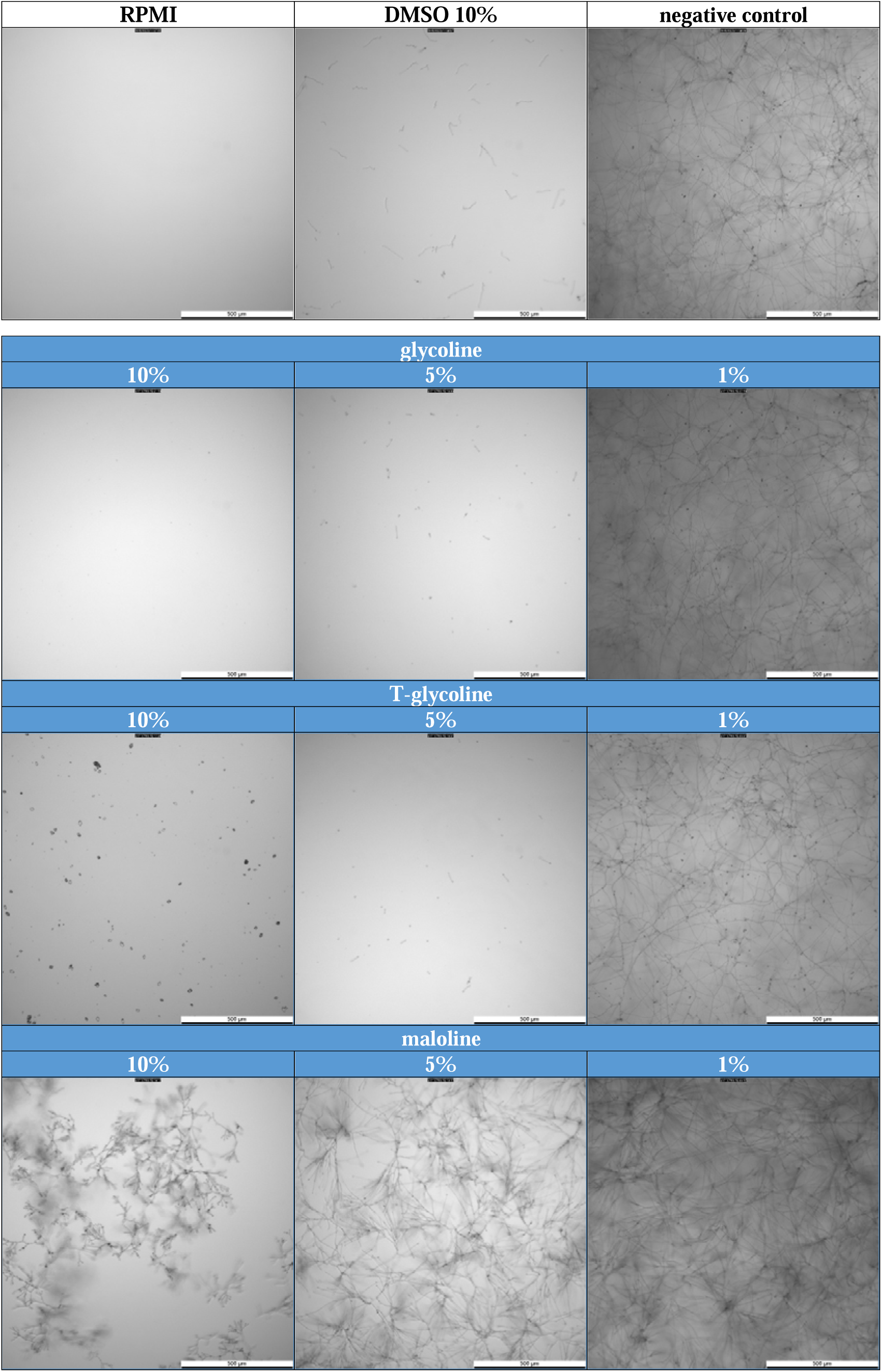
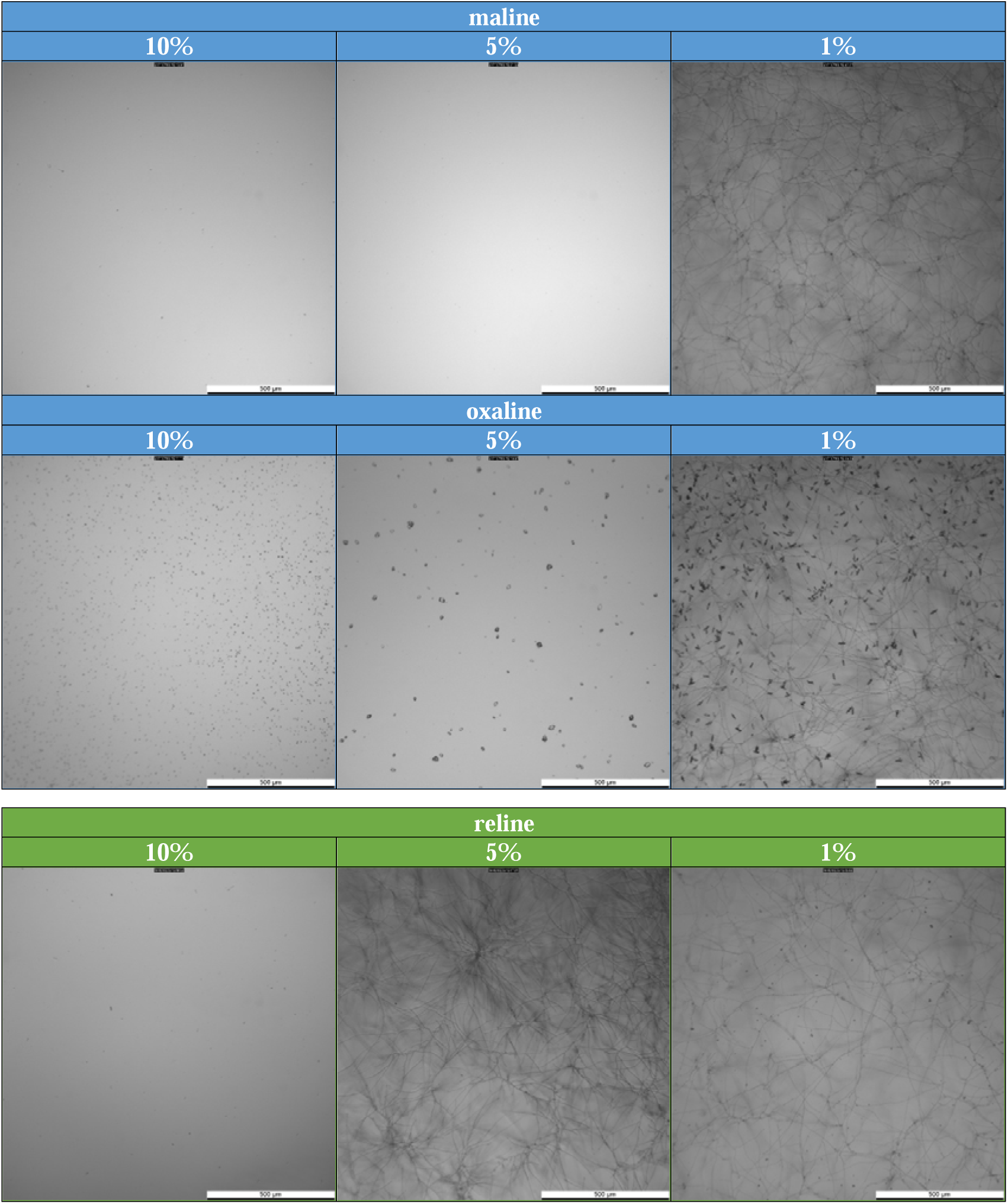
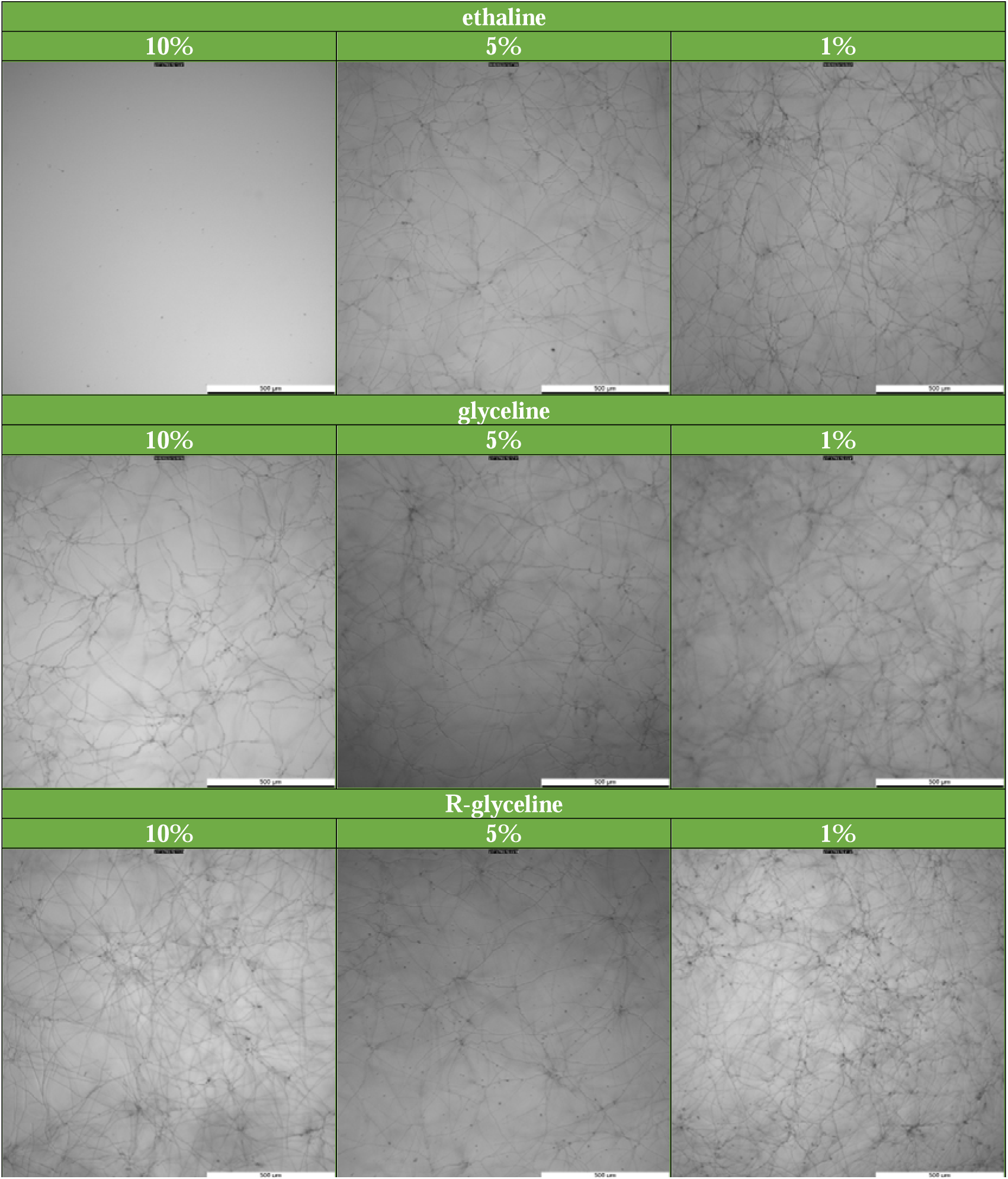
Optical microscopy images of *A. fumigatus* in the presence of the investigated DESs.

**Figure S10.**
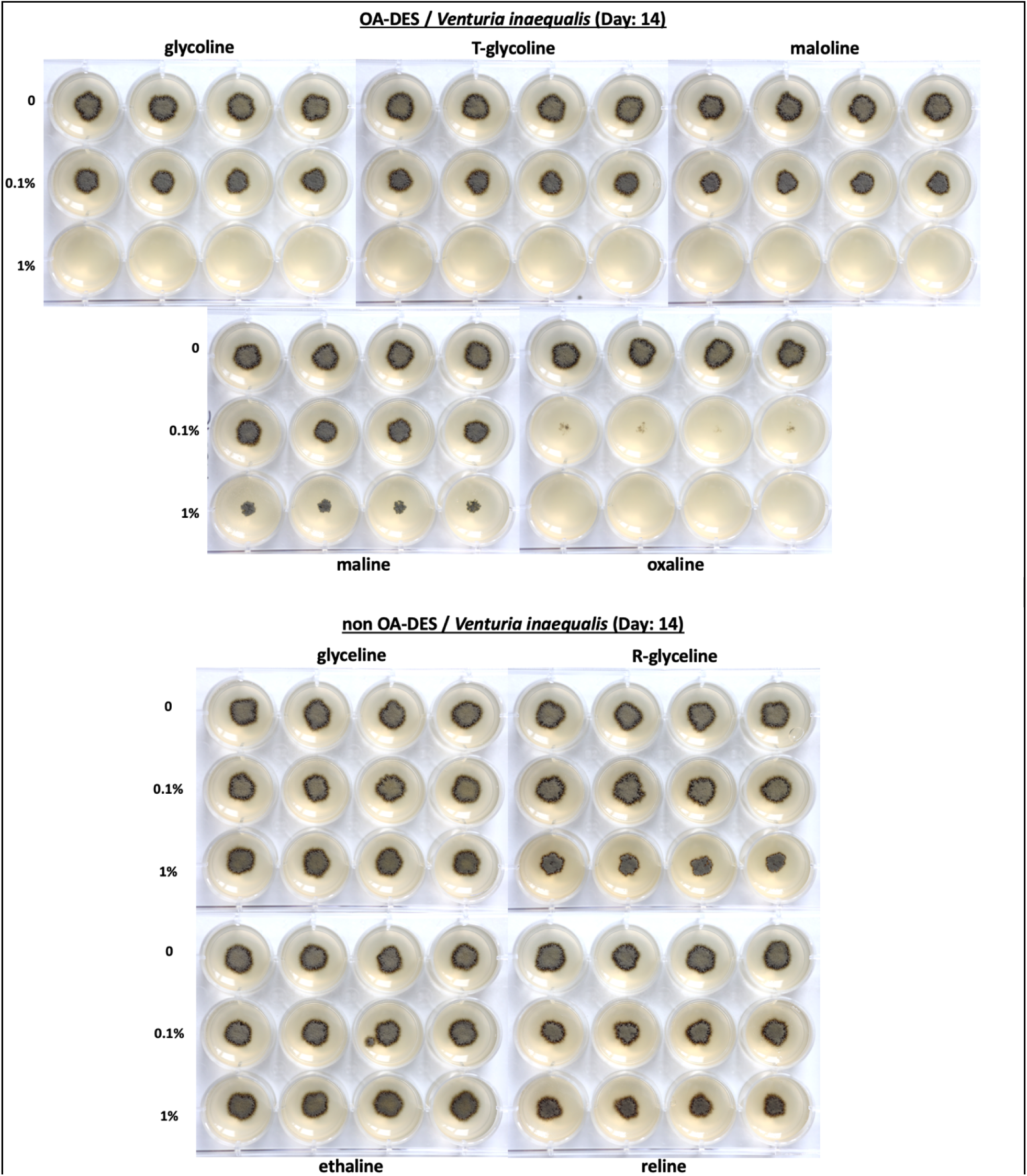
Images of the *V. inaequalis* treated with the investigated DESs.

**Figure S11.**
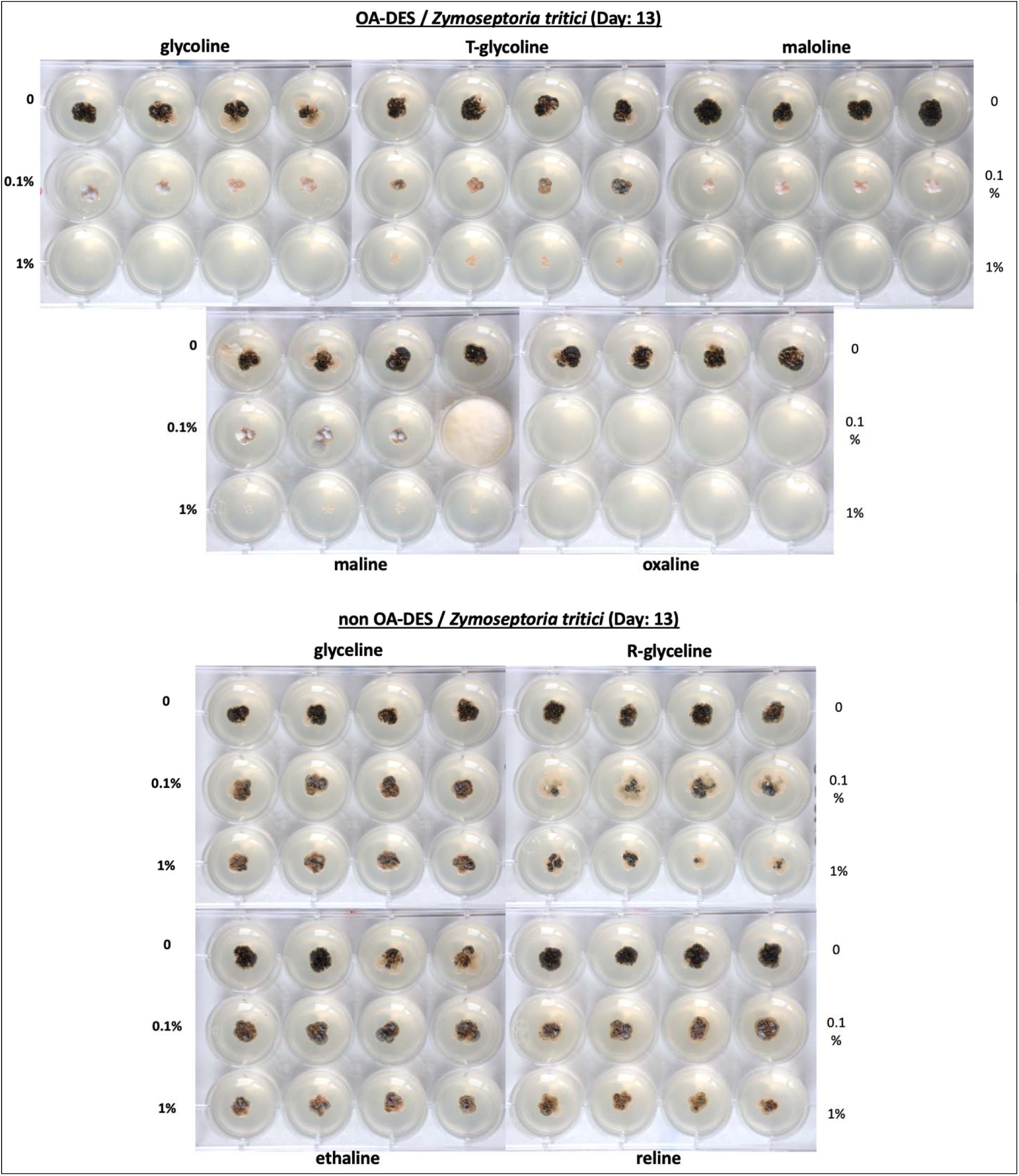
Images of the *Z. tricti* treated with the investigated DESs.

## Notes

### Competing Interest Statement

The authors have declared no competing interest.

### Summary of Updates

Reordered the authors to A. M. Kocot followed by M. Plotka

